# A discrete region of the D4Z4 is sufficient to initiate epigenetic silencing

**DOI:** 10.1101/2025.02.19.639175

**Authors:** Ellen M. Paatela, Faith G. St. Amant, Danielle C. Hamm, Sean R. Bennett, Taranjit S. Gujral, Silvère M. van der Maarel, Stephen J. Tapscott

## Abstract

The DUX4 transcription factor is briefly expressed in the early embryo and is epigenetically repressed in somatic tissues. Loss of epigenetic repression can result in the aberrant expression of DUX4 in skeletal muscle and can cause facioscapulohumeral dystrophy (FSHD). Multiple factors have been identified as necessary to maintain epigenetic silencing of *DUX4* in skeletal muscle, but whether specific sequences at the *DUX4* locus are sufficient for epigenetic silencing has been unknown. We cloned fragments of the D4Z4 macrosatellite repeat, the DNA region that encompasses the *DUX4* retrogene, adjacent to a reporter driven by a constitutive promoter and identified a single fragment sufficient to epigenetically repress reporter gene expression. Previously identified suppressors of *DUX4* expression—SETDB1, ATF7IP, SIN3A/B, and LRIF1—were necessary for silencing activity and p38 inhibitors enhanced suppression. These findings identify a key regulatory sequence for D4Z4 epigenetic repression and establish a model system for mechanistic and discovery studies.

## Introduction

Facioscapulohumeral dystrophy (FSHD) is the third most common muscular dystrophy, estimated to effect one in every 8-10,000 individuals (1), and is characterized by progressive wasting of muscles in the face and upper extremities (2,3). Current treatments for FSHD focus on alleviation of symptoms, as there is no cure (4). FSHD is caused by misexpression of the transcription factor double homeobox 4 (DUX4) in skeletal muscle (5). DUX4 is normally expressed as a short ‘burst’ in the cleavage stage embryo and at low levels in the testes and is epigenetically silenced in most other somatic tissues (5–7). Loss of this epigenetic repression can lead to DUX4 misexpression and activation of an embryonic program in skeletal muscle, eventually leading to FSHD pathogenesis (8–10).

The *DUX4* gene is located within each 3.3 kilobase (kb) tandem repeat unit of the D4Z4 macrosatellite arrays located in the subtelomeric regions of chromosomes 4q and 10q (11–15). There are two genetic causes of FSHD (16): FSHD1, which represents ∼95% of FSHD cases, is caused by a loss of D4Z4 repeat copy number from 8-100 to 1-10 copies (17,18), and FSHD2 (∼5% of cases) which is due to mutation of epigenetic regulators, such as Structural Maintenance of Chromosomes flexible Hinge Domain Containing 1 (SMCHD1) (>95% of FSHD2 cases), its binding partner Ligand Dependent Nuclear Receptor Interacting Factor 1 (LRIF1), or DNA methyltransferase 3 beta (DNMT3B) (19–21). Both FSHD1 and FSHD2 result in D4Z4 chromatin relaxation and DNA hypomethylation with the subsequent mis-expression of DUX4 in skeletal muscle.

While SMCHD1 binding to the DUX4 locus and the mouse Dux locus is implicated in the initial epigenetic silencing of DUX4 expression during development and establishment of DNA methylation (22,23), many factors have been identified that maintain epigenetic repression of DUX4 expression in somatic cells and skeletal muscle (8,24,25). A previous study in our lab identified many of these drivers by precipitation of D4Z4 repeats and unbiased proteomics analysis (26). Components of the Nucleosome Remodeling and Deacetylase (NuRD) complex together with SIN3 Transcriptional Regulator Family Members A and B (SIN3A and SIN3B) and Chromatin Assembly Factor-1 (CAF-1) complexes were identified as suppressors of *DUX4* expression (26). Histone H3 lysine 9 trimethylation (H3K9me3) loss at the D4Z4 is associated with FSHD (27) and loss of histone lysine methyltransferase (HKMT) SET Domain Bifurcated Histone Lysine Methyltransferase 1 (SETDB1) and Tripartite Motif Containing 28 (TRIM28), which complexes with HKMT Suppressor of Variegation 3-9 Homolog 1 (SUV39H1), led to increased *DUX4* expression in myoblasts (26,28). Several other chromatin-related D4Z4 suppressors have also been identified, including FSHD2 disease modifiers SMCHD1 and LRIF1, SMCHD1 interactor RuvB-like AAA ATPase 1 (RUVBL1), histone H3 lysine 4 demethylase Lysine Demethylase 1A (KDM1A), transcriptional repressor Yin Yang 1 (YY1), endogenous RNA interference (RNAi) pathway members Dicer 1 (DICER1) and Argonaut RISC catalytic component 2 (AGO2), and chromatin binding proteins Heterochromatin Protein 1 gamma (HP1γ), Cohesin, and CCCTC-binding factor (CTCF) (5,19,20,26,27,29–33). The large number of regulatory factors at the D4Z4 has made identification of the most critical silencing pathways difficult.

In this study, we designed a silencing reporter system to identify specific sequences in the D4Z4 that are sufficient for epigenetic silencing. Our silencing reporter system identified one discrete sequence in the D4Z4, termed D4Z4-S5, that confers epigenetic silencing activity and shows enrichment for SMCHD1 and LRIF1 binding. A candidate small interfering RNA (siRNA) panel of known D4Z4-regulatory proteins revealed several factors necessary to silence the D4Z4-S5 reporter, including SETDB1, its binding partner Activating Transcription Factor 7 Interacting Protein (ATF7IP), SIN3A/3B, and LRIF1. The D4Z4-S5 reporter was also further suppressed by p38 inhibitors, which are known to suppress DUX4 expression in FSHD skeletal muscle cells. Our study shows that initiation of epigenetic silencing, as measured by our reporter assay, is not distributed across the full D4Z4 unit but instead limited to one region and is regulated by specific factors. This reporter system also provides a simple and robust tool for identification of key D4Z4 regulators and potential therapeutics.

## Results

### Specific regions in the LRIF1 promoter and the D4Z4 suppress expression of an integrated reporter

To functionally assess the effect of specific D4Z4 sequences on silencing activity, we generated reporter constructs with the test sequence of interest placed upstream of a CMV promoter driving green fluorescence protein (*GFP*) expression (Fig. 1A) and used homologous recombination to introduce these reporter constructs into the *AAVS1* (*PPP1R12C*) locus in HeLa cells. The AAVS1 locus is considered a “safe harbor” due to its low intrinsic silencing and no discernible phenotype upon perturbation (34). HeLa cells were used to avoid the dynamic regulation of D4Z4 expression during myogenic differentiation (35,36).

**Figure 1:**
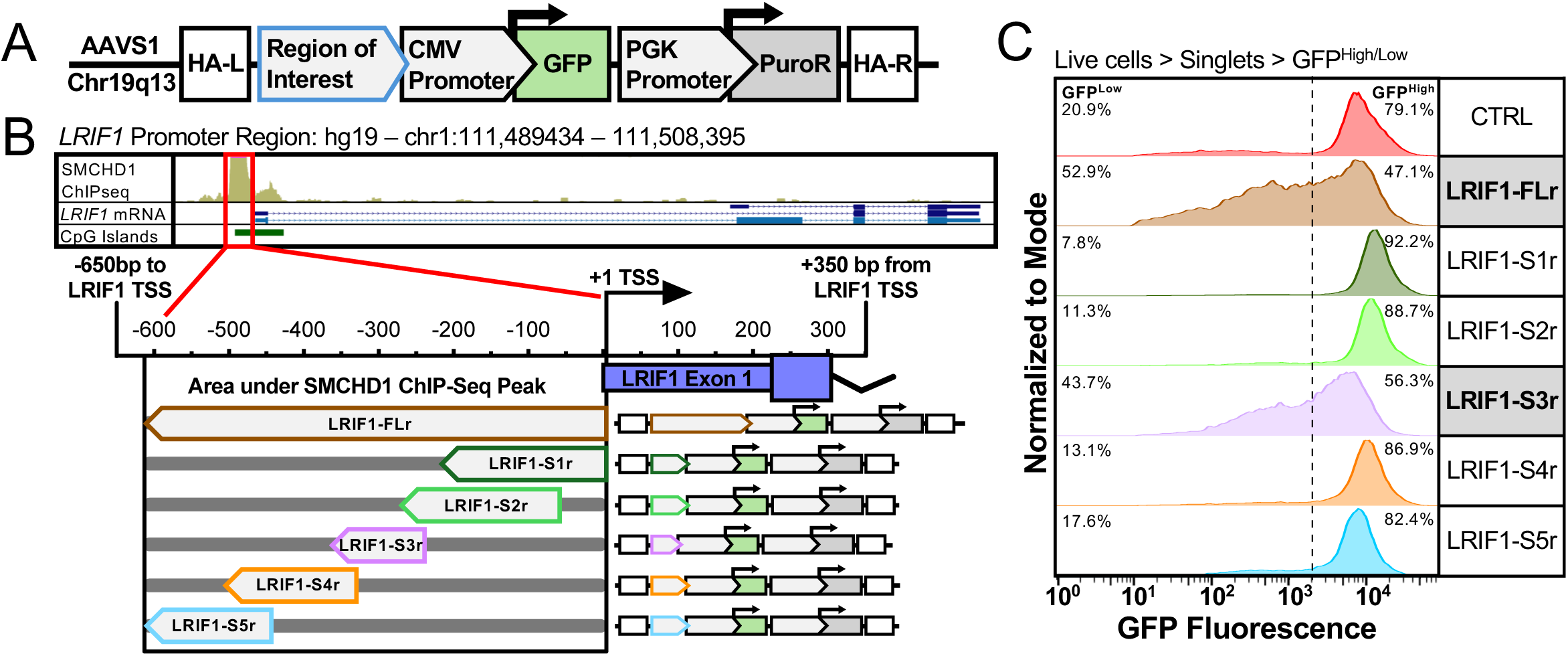
A safe harbor system identifies silencing activity driven by specific sequences in the *LRIF1* promoter. **(A)** Schematic of the safe harbor construct integrated into the *AAVS1* locus in HeLa cells. A sequence of interest is inserted upstream of a constitutive CMV promoter driving GFP. **(B)** Top: UCSC genome browser track of SMCHD1 ChIP-seq peaks (37), LRIF1 mRNA, and CpG islands at the *LRIF1* promoter region. Bottom: Sequence maps of LRIF1 promoter region segments. Arrowheads indicate orientation of sequence relative to genomic sequence and construct schematics indicate orientation of sequences relative to integrated reporter constructs. **(C)** Flow cytometry histogram of GFP fluorescence in cells (n=1 biological replicate) containing various integrated *LRIF1* promoter silencing constructs. Histograms are normalized to the mode. Percentages of cell populations classified GFP^High^ or GFP^Low^ are indicated.

Amongst D4Z4 suppressors, SMCHD1 has an established role in initiation and maintenance of D4Z4 epigenetic repression. Previous ChIP analysis established that SMCHD1 and LRIF1 bind a discrete region adjacent to the *LRIF1* promoter and suppress *LRIF1* expression (37,38). Therefore, as a positive control for silencing activity, we inserted the full-length 617 nucleotide (nt) region containing the SMCHD1 ChIP-seq peak (termed LRIF1-FL) upstream of the CMV promoter into the reporter construct (Fig. 1B) and introduced this into the HeLa AAVS1 locus. Because this region is directly upstream of the *LRIF1* transcriptional start site (TSS), we reversed the orientation of the LRIF1-FL sequence (LRIF1-FLr) to avoid changes in GFP expression due to transcription from the LRIF1 promoter. The directionality of integration within the AAVS1 construct is shown in Fig. 1B. For comparison, we also introduced a 200 nt region (CTRL, chr11:114,494,095-114,494,295), previously identified as a sequence with no regulatory activity (39), as a control and likely neutral sequence. Cells were expanded under selection as a polyclone and accurate integration was confirmed by polymerase chain reaction (PCR) genotyping. To ensure low GFP levels were due to silencing and not a stable GFP^Low^ population, GFP^High^ cells were sorted via fluorescence activated cell sorting (FACS) and cultured for two weeks to allow for construct silencing. Flow cytometry showed that nearly 80% of the CTRL cells expressed GFP at relatively high levels, whereas cells with LRIF1-FLr had approximately 50% of cells showing low GFP fluorescence, approaching a bimodal distribution (Fig. 1C). Reversing the orientation of LRIF1-FL was necessary for silencing, as flow cytometry of the LRIF1-FL forward sequence (LRIF1-FLf) did not induce silencing, indicating that the presence of the *LRIF1* promoter sequence directly upstream of the TSS was likely driving transcriptional activation (Supplementary Material, Fig. S1A-B). Fragmentation of the LRIF1-FLr into five overlapping segments showed silencing activity above control levels and was restricted to the 112 nt LRIF1-S3r subfragment (Fig. 1C) centered under the SMCHD1/LRIF1 ChIP peaks (38) (Fig. 1B), providing a positive control to show that the integrated reporter can be silenced by a region shown to bind SMCHD1/LRIF1. To ensure that directionality was not a specific driver of silencing activity, the reverse compliment of LRIF1-S3, LRIF1-S3f, which contains the sequence in the correct orientation relative to the endogenous *LRIF1* TSS, was integrated into the AAVS1 system (Supplementary Material, Fig. S1A). Regardless of orientation, LRIF1-S3 showed GFP suppression compared to the control sequence, indicating this sequence is truly a silencer regulatory element (Supplementary Material, Fig. S1B).

To distinguish whether D4Z4 silencing is localized to a specific region of the D4Z4 or is broadly distributed, the 3.3kb D4Z4 repeat unit was segmented into 14 overlapping fragments, cloned into the reporter construct, and integrated at the AAVS1 locus (Fig. 2A). Flow cytometry revealed that over 40% of cells showed substantial GFP suppression with a single 401 nt region, D4Z4 Segment 5 (D4Z4-S5), with minimal silencing activity in other D4Z4 regions (Fig. 2B). D4Z4-S5 was the only region with a greater percentage of GFP^Low^ cells than the CTRL sequence. Interestingly, this GFP silencing activity was orientation dependent, as reversing the sequence (D4ZR-S5r) did not show increased GFP suppression compared to the CTRL reporter (Supplementary Material, Fig. S2A-B). To ensure that this bimodal distribution of GFP fluorescence in D4Z4-S5 was due to *de novo* AAVS1 silencing and not due to a stable population of GFP^Low^ cells in the polyclonal reporter line, cells containing the CTRL reporter, or cell lines with D4Z4 reporter constructs were sorted via FACS for GFP^High^ cells and maintained in culture. At 1- and 5-weeks post sort, flow cytometry was performed and showed substantial GFP suppression at week 5 relative to control levels only in D4Z4-S5, confirming that the GFP suppression is not due to expression artifacts in a polyclonal population of cells, but by sequence-dependent silencing over time (Supplementary Material, Fig. S2C-D). Further fragmentation of D4Z4-S5 (Fig. 2C) localized the repressive region to a 146 nt fragment (D4Z4-S5.4, Fig. 2C-D), which showed a larger population of GFP^Low^ cells than the control.

**Figure 2:**
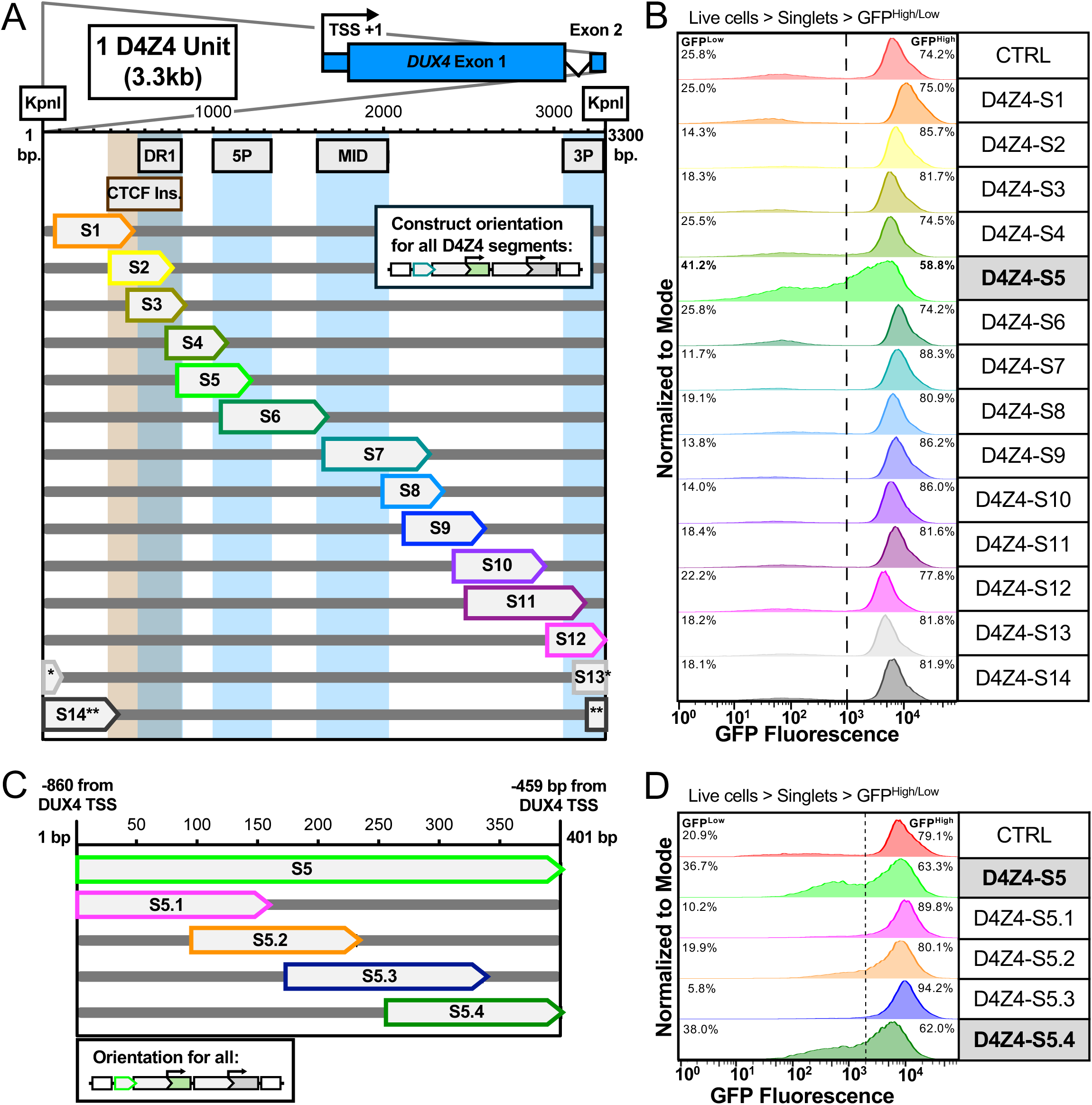
Segmentation of the D4Z4 reveals one sequence with silencing activity. **(A)** Schematic of one D4Z4 repeat unit and sequence segments inserted into the silencing reporter construct. Asterisks on D4Z4-S13 and D4Z4-S14 indicate segments span across the KpnI site into the next D4Z4 repeat unit. **(B)** Flow cytometry histogram of GFP fluorescence in cells containing various integrated D4Z4 segment constructs (n=1 biological replicate). Histograms are normalized to the mode. Percentages of cell populations classified GFP^High^ or GFP^Low^ are indicated. **(C)** Schematic of D4Z4-S5 and sequence sub-fragments inserted into the silencing reporter construct. **(D)** Flow cytometry histogram of GFP fluorescence in cells (n=1 biological replicate) containing various integrated D4Z4-S5 subfragment silencing constructs. Histograms are normalized to the mode. Percentages of cell populations classified GFP^High^ or GFP^Low^ are indicated.

GFP silencing regions D4Z4-S5, D4Z4-S5.4, LRIF1-FL, and LRIF1-S3 have similar GC content and percentage of CpG dinucleotides to other regions of the D4Z4 (Supplementary Material, Table S1), suggesting that other sequence-specific aspects may account for silencing activity. LRIF1-S3 contains two regions with high conservation in placental mammals predicted by PhyloP (40) (Supplementary Material, Fig. S3A); a minimal subregion (LRIF1-S3f Minimal Sufficiency), containing these motifs is sufficient for GFP silencing, whereas deletion of one of the regions (LRIF1-S3fΔsite#1) appears to reduce GFP silencing (Supplementary Material, Fig S3B). These results indicate site#1 is important for GFP silencing activity, though motif analysis did not reveal any obvious regulator candidates in either conserved site. In contrast to LRIF1-S3, PhyloP predicted no conserved motifs among 100 representative vertebrates in D4Z4-S5.4, and fractionation to smaller fragments did not strongly map silencing activity to a specific smaller region (Supplementary Material, Fig S3C-D).

### Epigenetic repression of D4Z4-S5 contributes to silencing activity

The D4Z4-S5 region partially overlaps with the previously described DR1 and 5P regions of the D4Z4 (41,42), and D4Z4-S5.4 is centered in the 5P region. Previous studies have shown that both regions have relatively high percentage CpG methylation and are specifically hypomethylated in FSHD (41–44). The D4Z4 also shows high levels of repressive histone modifications, where a loss of these modifications is associated with DUX4 expression and FSHD pathogenesis (27,36,38,45). To determine the contributions of DNA methylation and repressive histone modifications to GFP silencing in our AAVS1 reporter system, D4Z4-S5 and LRIF1-FLr cells were treated with the DNA demethylating agent 5-aza-2’-deoxycytidine (5-aza-dC) or the histone deacetylase inhibitor entinostat for three days, and GFP fluorescence levels were quantified via flow cytometry at several timepoints after treatment (Fig. 3A). Treatment with either drug increased the proportion of GFP^High^ cells in the population, indicating a loss of GFP suppressive mechanisms (Fig. 3B-C). Increased time in culture also revealed a re-emergence of the GFP^Low^ population and a restoration of pre-treatment median GFP fluorescence levels at days 7 and 10 (Fig. 3B-C, right). These results suggest the silencing in D4Z4-S5 and LRIF1-FLr is due, in part, to epigenetic mechanisms.

**Figure 3:**
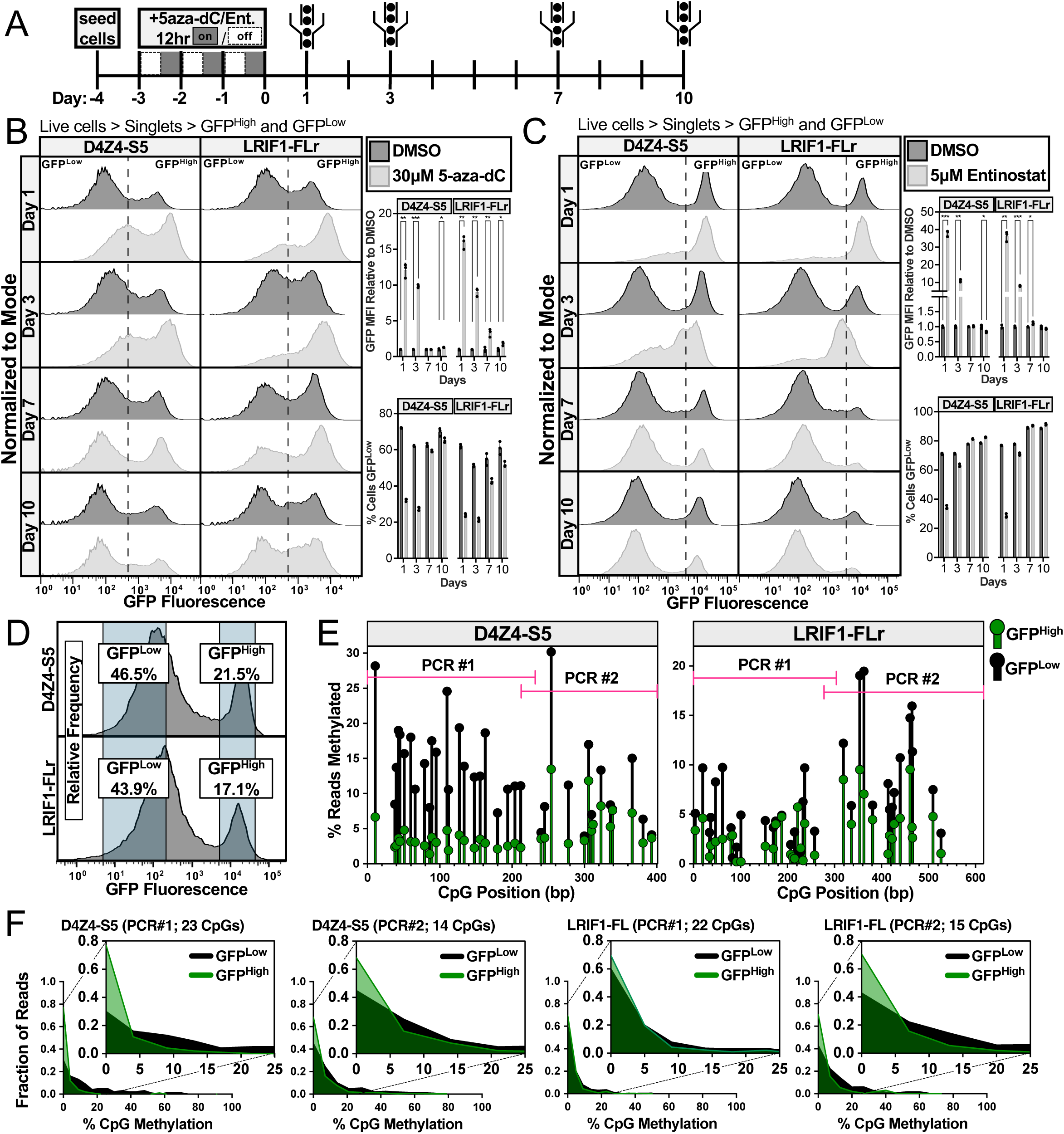
GFP silencing activity of D4Z4-S5 is partially due to *de novo* DNA methylation and repressive histone modifications. **(A)** Schematic of experimental time course. Cells were treated with 5-aza-dC or Entinostat for 3 days with a 12 hour on/off schedule and harvested for flow cytometry analysis at 1, 3, 7, and 10 days after washout. **(B-C)** Left: Representative singleton GFP fluorescence histograms of CTRL, D4Z4-S5 and LRIF1-FLr cells at various timepoints after 5-aza-dC **(B)** or entinostat **(C)** treatment. Histograms are normalized to the mode of the population. Right Top: Fold change in GFP MFI at various timepoints after treatment compared to the DMSO control. Data represent mean ± SD of biological replicates, n=3. Statistical significance was determined by Welch’s t-test: *p<0.05, **p<0.01, ***p<0.001. Right bottom: Percentage of GFP^Low^ cells in the population at various time points after treatment. Data represent mean ± SD of biological replicates, n=3. **(D)** GFP fluorescence histograms of D4Z4-S5 and LRIF1-FLr, plotted based on relative frequency. Blue boxes indicate FACS sorting gates for GFP^High^ and GFP^Low^ populations used in subsequent DNA methylation analysis. **(E)** Lollipop plots of methylation levels at individual CpGs for D4Z4-S5 (left) and LRIF1-FLr (right). The percentage of nanopore sequencing reads of bisulfite PCR products with methylation (no C>T conversion) at each indicated CpG is plotted for GFP^Low^ (black) and GFP^High^ (green) populations. **(F)** Fraction of bisulfite PCR nanopore sequencing whole reads with various percentages of CpG methylation for GFP^Low^ (black) and GFP^High^ (green) cell populations. Inset graph shows read fractions with less than 25% total CpG methylation. Raw bisulfite PCR sequencing reads in **Supplemental Table 2**.

To determine whether GFP suppression is correlated with higher DNA methylation, D4Z4-S5 and LRIF1-FLr cells were sorted into GFP^High^ and GFP^Low^ populations (Fig. 3D), genomic DNA was bisulfite converted, AAVS1 reporter construct regions were PCR-amplified, and amplicons were sequenced using long-read nanopore technology to determine methylation status (Supplementary Material, Table S2). Compiling two PCR amplicons for each, higher methylation in GFP^Low^ cells was distributed throughout most CpG sites in D4Z4-S5 (Fig. 3E, left) and larger differences in CpG methylation between GFP^High^ and GFP^Low^ populations were concentrated in one region, between 300 and 500nt, of LRIF1-FLr (Fig. 3E, right). AAVS1 PCR amplicons from both D4Z4-S5 and LRIF1-FLr showed higher levels of overall CpG methylation per read in the GFP^Low^ cells compared to the GFP^High^ cells (Fig. 3F). In the GFP^Low^ cells, less than half of the reads are completely unmethylated in both D4Z4-S5 amplicons and LRIF1-FLr PCR#2 (in LRIF1-FLr PCR#1, less than 60% of reads are unmethylated), while in the GFP^High^ populations, 60-80% of reads are completely unmethylated in all AAVS1 amplicons, correlating unmethylated AAVS1 DNA with higher GFP fluorescence (Fig. 3F). The presence of unmethylated reads in the GFP^Low^ cell populations, however, indicates that DNA methylation is only partially responsible for GFP silencing, and suggests that other epigenetic mechanisms also contribute to sequence-dependent GFP suppression in D4Z4-S5 and LRIF1-FLr.

### Histone methyltransferase and histone de-acetylase activity contributes to silencing at D4Z4-S5 and LRIF1-FLr

Previously, we showed that specific histone methyltransferases, de-acetylases, and components of the NuRD complex were necessary to maintain D4Z4 epigenetic repression in human myoblasts (26). To determine if these known D4Z4 regulators are also responsible for sequence-dependent silencing in D4Z4-S5 and LRIF1-FLr reporter constructs, a focused siRNA-mediated knockdown (KD) panel of factors known to epigenetically regulate D4Z4 epigenetic repression (26) was performed in reporter cells. Loss of CHD4 and MBD1 (Supplementary Material, Fig. S4A-D), components of the NuRD complex, did not strongly de-repress GFP expression in the D4Z4-S5 or LRIF1-FLr constructs, though loss of MBD1 led to a small but significant increase in the average GFP median fluorescence intensity (MFI) relative to siCTRL. However, loss of the H3K9 histone methyltransferase SETDB1 robustly de-repressed GFP expression in both D4Z4-S5 and LRIF1-FLr when compared to CTRL cells (Fig. 4A, Supplementary Material, Fig. S5A). The relative GFP MFI increased roughly 4- and 7-fold in D4Z4-S5 and LRIF1-FLr cells, respectively, while decreasing in CTRL cells (Fig. 4A, right). Loss of ATF7IP, a binding partner of SETDB1 that increases SETDB1 stability and methyltransferase activity (46), significantly increased GFP fluorescence in all reporter cell lines; however, the fold change in relative MFI is an order of magnitude greater for D4Z4-S5 and LRIF1-FLr cells relative to CTRL (Fig. 4B, Supplementary Material, Fig. S5D). ChIP-quantitative PCR (qPCR) analysis revealed that loss of SETDB1 or ATF7IP lead to a significant loss of H3K9me3 specifically at AAVS1 D4Z4-S5 and LRIF1-FLr inserts, whereas the endogenous loci and control regions (a gene desert region (GDR) that has high baseline H3K9me3 and a promoter region of housekeeping gene ribosomal protein L13a (RPL13A) that has low baseline H3K9me3) did not show a change in H3K9me3 levels (Fig. 4C, Supplementary Material, Fig. S5E). Of interest, knockdown of sequence-specific SETDB1/ATF7IP recruitment factor TRIM28 did not de-repress these regions (Supplementary Material, Fig. S4E-F) suggesting alternative mechanisms are required for recruitment and/or silencing activity of SETDB1 and ATF7IP at D4Z4-S5 and LRIF1-FLr.

**Figure 4:**
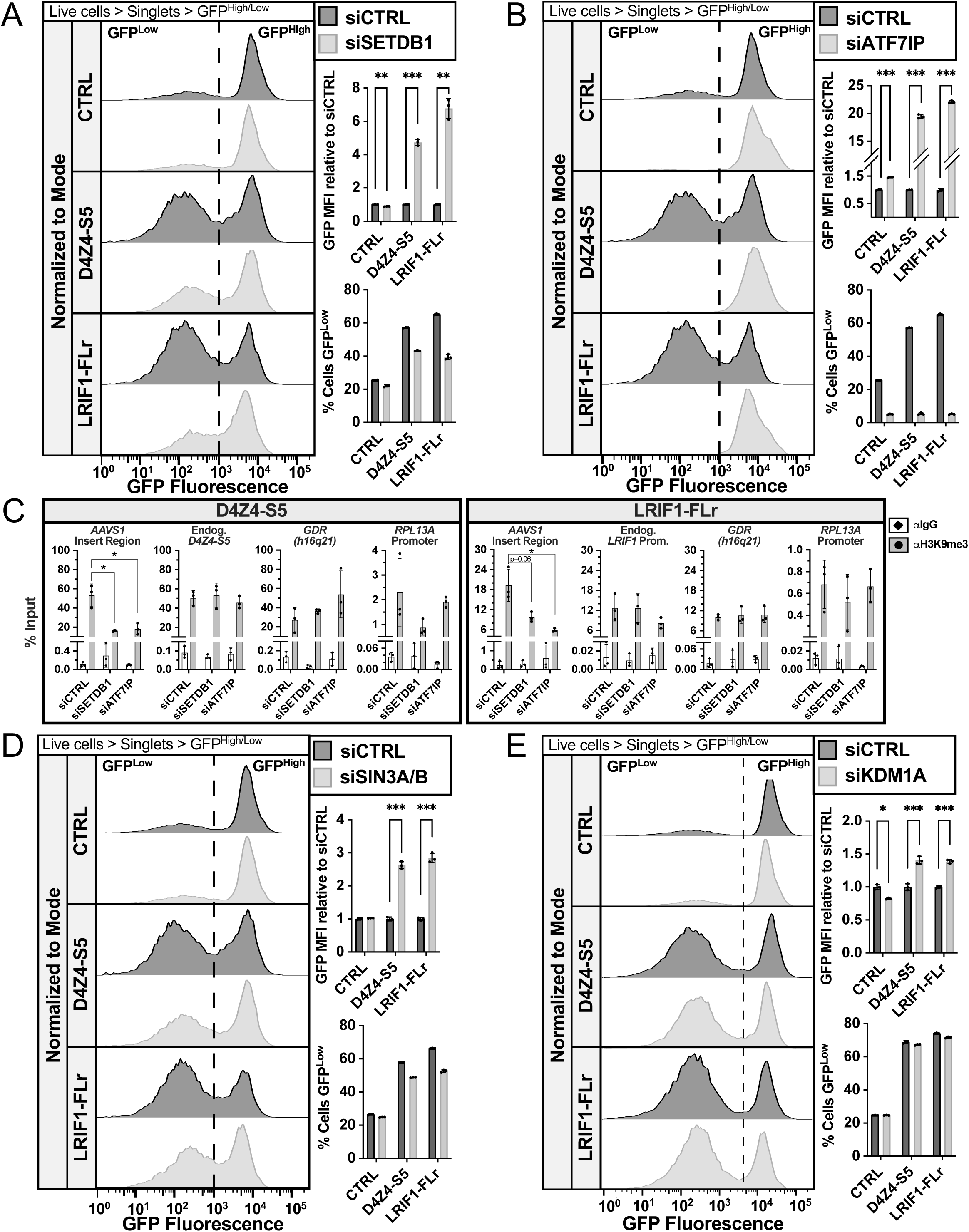
Loss of histone methyltransferases and deacetylases specifically relieves D4Z4-S5 and LRIF1-FLr GFP silencing. **(A-B, D-E)** Left: Representative singleton GFP fluorescence histograms of CTRL, D4Z4-S5 and LRIF1-FLr cells treated with CTRL or SETDB1 **(A)**, ATF7IP **(B)**, SIN3A/B **(D)**, or KDM1A **(E)** siRNAs. Histograms are normalized to the mode of the population. Right Top: Fold change in GFP MFI upon siRNA-mediated knockdown of SETDB1 **(A)**, ATF7IP **(B)**, SIN3A/B **(D)**, or KDM1A **(E)** compared to siCTRL. Data represent mean ± SD of biological replicates, n=3. Statistical significance was determined by Welch’s t-test: *p<0.05, **p<0.01, ***p<0.001. Right bottom: Percentage of GFP^Low^ cells in the population at various time points after siRNA treatment. Data represent mean ± SD of biological replicates, n=3. **(C)** ChIP-qPCR of H3K9me3 occupancy at various loci in D4Z4-S5 and LRIF1-FLr cells upon treatment with siCTRL, siSETDB1, or siATF7IP. Data is graphed as the percent of total chromatin input that was immunoprecipitated by IgG or H3K9me3 antibodies. Data represent mean ± SD of biological replicates, n=3. Statistical significance was determined by Welch’s t-test: *p<0.05.

In addition to H3K9 methyltransferases, loss of other repressive histone modifying enzymes, such as histone deacetylases SIN3A and SIN3B and H3K4 demethylase KDM1A, were previously found to relieve suppression of DUX4 in myoblasts (26). Knockdown of SIN3A and SIN3B in our AAVS1 reporter cells showed a 2.5-fold increase in GFP fluorescence in D4Z4-S5 and LRIF1-FLr cells and not in CTRL cells, indicating histone deacetylation may also contribute to silencing (Fig. 4D, Supplementary Material, Fig. S5C). Interestingly, knockdown of KDM1A did not show a substantial increase in GFP fluorescence, with changes in relative GFP MFI less than 1.5-fold (Fig. 4E, Supplementary Material, Fig. S5D), indicating demethylation of histone H3 on lysine 4 (H3K4) is likely not a driver of sequence-specific GFP suppression in D4Z4-S5 and LRIF1-FLr.

### SMCHD1 and LRIF1 are enriched at D4Z4-S5 and LRIF1-FLr yet display different silencing roles

Prior studies from our lab and others document SMCHD1/LRIF1 binding to the endogenous region corresponding to LRIF1-FLr (37,38,47). ChIP-qPCR analysis of our silencing reporter cells confirmed SMCHD1 and LRIF1 binding to the endogenous *LRIF1* promoter region and identified enriched binding at both the LRIF1-FLr and D4Z4-S5 transgene AAVS1 inserts (Fig. 5A). Of additional interest, ChIP-qPCR also identified enrichment of these factors at the endogenous D4Z4-S5.4 region (Fig. 5A). Knockdown of LRIF1 (short and long isoforms) increased reporter expression in both the LRIF1-FLr and D4Z4-S5 cells with a significant increase in GFP MFI (Fig. 5B, Supplementary Material, Fig. S6A-C), supporting our ChIP data showing specific LRIF1 binding and silencer recruitment activity. In contrast, knockdown of SMCHD1 shows a significant decrease in reporter expression for both cell lines, indicating a potential positive regulatory role for reporter expression (Fig. 5C, Supplementary Material, Fig. S6A-C). Although SMCHD1 can have a role as a transcriptional co-activator in some contexts (48), SMCHD1 is known to negatively regulate *LRIF1* expression (38). GFP suppression upon loss of SMCHD1 can potentially be explained by increased levels of the silencer LRIF1; however, differences in LRIF1 RNA and protein levels were not significant (Supplementary Material, Fig. S6A-C). In summation, LRIF1 and SMCHD1 are recruited in a sequence-specific manner to D4Z4-S5 and LRIF1-FLr, though only LRIF1 seems to contribute to silencing activity in our assays.

**Figure 5:**
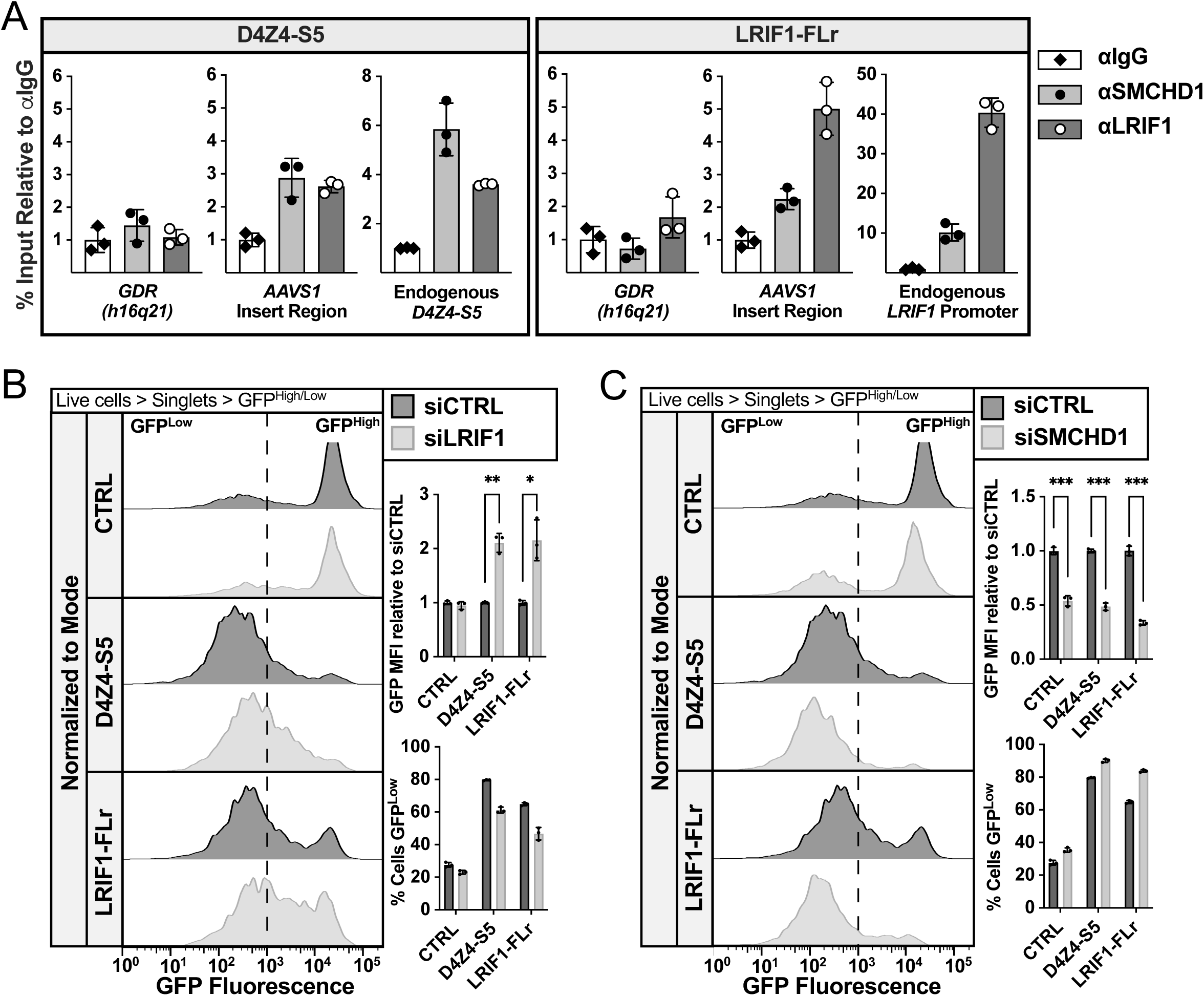
SMCHD1 and LRIF1 are enriched at D4Z4-S5 and LRIF1-FLr yet display different silencing roles. **(A)** ChIP-qPCR analysis of SMCHD1 and LRIF1 occupancy at various endogenous loci and the AAVS1 silencing construct transgene in D4Z4-S5 and LRIF1-FLr cells. Data represents the % of input chromatin immunoprecipitated by SMCHD1 and LRIF1 antibodies relative to IgG of three technical triplicates (n=1 biological replicate). **(B-C)** Left: Representative singleton GFP fluorescence histograms of CTRL, D4Z4-S5 and LRIF1-FLr cells treated with CTRL or LRIF1 **(B)** or SMCHD1 **(C)** siRNAs. Histograms are normalized to the mode of the population. Right Top: Fold change in GFP MFI upon siRNA-mediated knockdown of LRIF1 **(B)** or SMCHD1 **(C)** compared to siCTRL. Data represent mean ± SD of biological replicates, n=3. Statistical significance was determined by Welch’s t-test: *p<0.05, **p<0.01, ***p<0.001. Right bottom: Percentage of GFP^Low^ cells in the population at various time points after siRNA treatment. Data represent mean ± SD of biological replicates, n=3.

### D4Z4-S5 and LRIF1-FLr cell lines are sensitive to p38 pathway inhibition and can be used to screen for novel FSHD therapeutics

Several studies have identified p38 inhibitors, beta-2-adrenergic receptor agonists, bromodomain and extraterminal (BET) inhibitors, and G-quadruplex stabilizers as repressors of *DUX4* expression, with several of these drugs advancing to clinical trials for treatment of FSHD (49–55). To assess the efficacy of our AAVS1 silencing reporter system as a tool for screening FSHD therapeutics, CTRL, LRIF1-FLr, and D4Z4-S5 cells were treated with several candidate drugs at concentrations sufficient to suppress *DUX4* and DUX4 target genes in FSHD myoblasts. Compared to dimethyl sulfoxide (DMSO) treatment, p38 inhibitors losmapimod and SB203580 substantially suppressed GFP expression in a dose-dependent manner in D4Z4-S5, and to a lesser degree LRIF1-FLr, cells when compared to CTRL cells (Fig. 6A-B). Higher doses of losmapimod and SB203580 significantly decreased the median GFP fluorescence intensity and increased the proportion of GFP^Low^ cells in D4Z4-S5, while CTRL and LRIF1-FLr cells showed similar trends but to a lesser degree (Fig. 6C-D). These results indicate that p38 pathway inhibition may reduce DUX4 expression through interactions with the D4Z4-S5 sequence.

**Figure 6:**
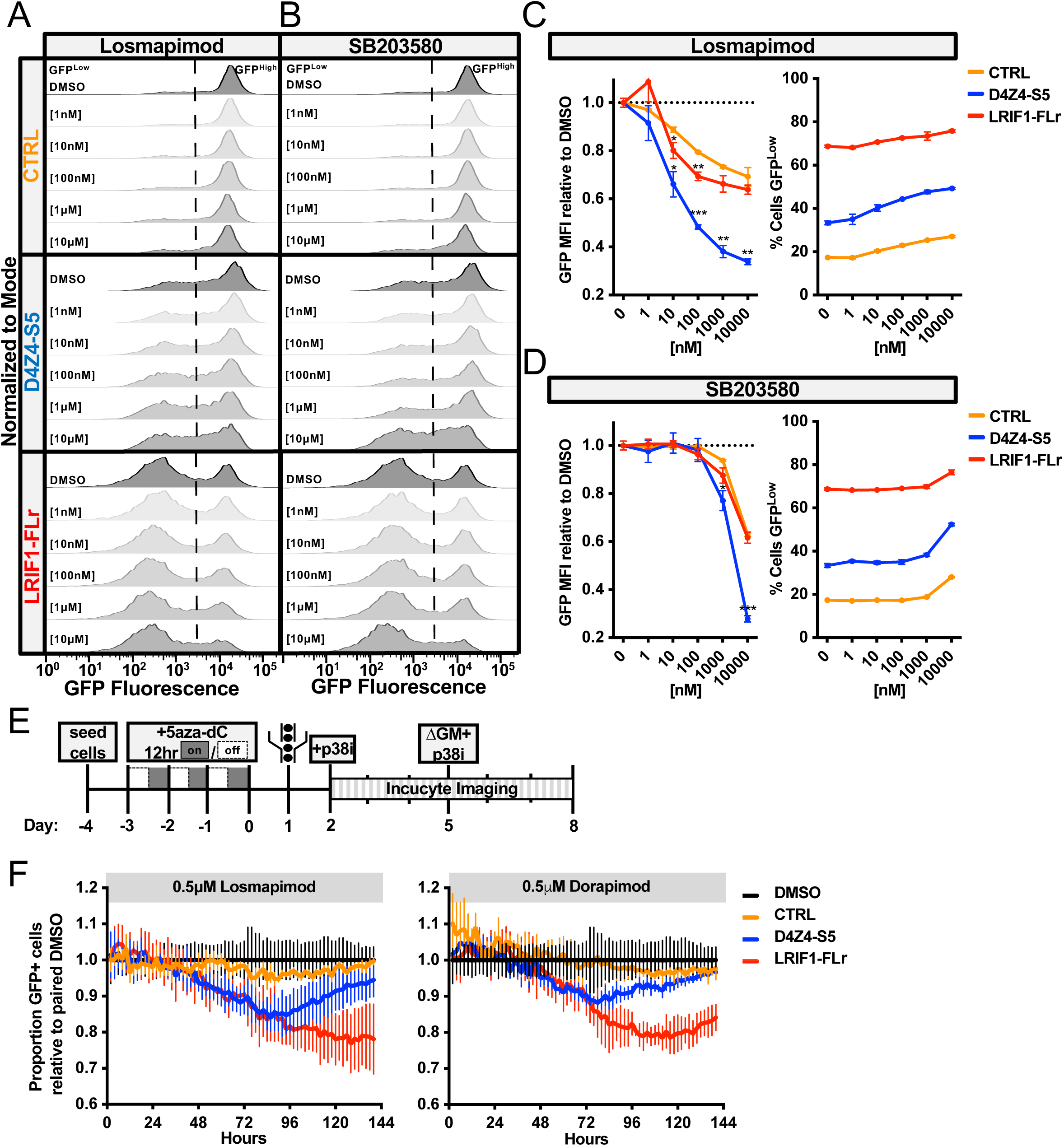
Candidate p38 inhibitor FSHD therapeutics inhibit GFP expression specifically in D4Z4-S5 and LRIF1-FLr. **(A-B)** Representative singleton GFP fluorescence histograms of CTRL, D4Z4-S5 and LRIF1-FLr cells treated for 3 days with DMSO or escalating dosages of p38 pathway inhibitors Losmapimod **(A)** or SB203580 **(B)**. Histograms are normalized to the mode of the population. **(C-D)** Left: Fold change in GFP MFI upon 3-day treatment with escalating dosages of Losmapimod **(C)**, or SB203580 **(D)** compared to DMSO treatment in CTRL (orange), D4Z4-S5 (blue), and LRIF1-FLr (red) cells. Data points represent mean fold change ± SD of biological replicates, n=3. Statistical significance was determined by Welch’s t-test: *p<0.05, **p<0.01, ***p<0.001. Right: Percentage of GFP^Low^ cells in the population upon treatment with escalating dosages of Losmapimod **(C)** or SB203580 **(D)**. Data represent mean ± SD of biological replicates, n=3. **(E)** Schematic of live cell imaging experimental time course. Cells were treated with 5-aza-dC for three days with a 12 hour on/off schedule. 24 hours after washout, GFP^High^ cells were FACS sorted into multi-well plates. 24 hours after sort, cells were treated with p38 inhibitors and incubated for 6 days in an Incucyte live cell imaging system, and images of each well were captured every 2 hours for GFP fluorescence quantification. On day 3 of Incucyte incubation, cell growth media was refreshed and p38 inhibitors were replenished. **(F)** Plots of GFP+ cell confluency over 6 days upon treatment with Losmapimod (left) or Dorapimod (right). Data shown is the GFP+ cell confluency in p38 inhibitor-treated wells relative to DMSO-treated (black) wells in CTRL (orange), D4Z4-S5 (blue), or LRIF1-FLr (red) cells. Data points are the average relative GFP+ cell confluency ± SD of biological replicate wells, n=6.

In contrast, beta-2-adrenergic receptor agonists clenbuterol and formoterol did not change levels of GFP expression in any of the cell lines (Supplementary Material, Fig. S7A-D). Treatment trends showed increases in median GFP fluorescence levels and decreased proportions of GFP^Low^ cells at higher doses, indicating that these drugs likely do not impact DUX4 expression through regulation of the D4Z4-S5 sequence. Treatment with BET agonist (+)- JQ1 also did not substantially alter levels of GFP fluorescence, though the highest dose had a significant suppressive effect on D4Z4-S5 and LRIF1-FLr (Supplementary Material, Fig. S7E-F). Surprisingly, treatment with G-quadruplex stabilizer berberine strongly suppressed GFP in a dose dependent manner specifically in D4Z4-S5 cells (Supplementary Material, Fig. S7G-H), despite the lack of known D4Z4 G-quadruplex ligands in the D4Z4-S5 sequence (53). Further study is necessary to determine the mechanism of berberine-mediated transcriptional silencing through D4Z4-S5.

Finally, to assess the efficacy of D4Z4-S5 as a platform for large scale therapeutic screens, GFP fluorescence changes in our AAVS1 reporter cells were assessed in a live-cell imaging system. Cells were treated with 5-aza-dC to ensure strong GFP expression before GFP^High^ cells were sorted via FACS into multi-well plates. These cells were then cultured in the presence of p38 inhibitors losmapimod or dorapimod for multiple days, during which live cell imaging quantified GFP+ cell confluency per well every 2 hours (Fig. 6E). p38-inhibitor treatment continuously suppressed LRIF1-FLr GFP expression over a six-day period, whereas D4Z4-S5 showed transient suppression maximal at 3.5 days with substantial recovery during longer treatment (Fig. 6F). Together these data indicate that LRIF1-FLr and D4Z4-S5 repressive activity is modulated by p38 signaling and that these AAVS1 reporter cells provide a sensitive platform for live-cell imaging-based assays to identify additional targets and compounds that modulate expression.

## Discussion

Although many prior studies have described factors necessary for epigenetic silencing of the D4Z4 repeats in somatic cells, it was unknown whether DNA sequences sufficient to initiate silencing were broadly distributed across the 3.3kb D4Z4 unit, or whether multiple copies of the D4Z4 array were necessary for silencing. The major finding of our current study is that, from the entire 3.3 kb D4Z4, only one discrete region of ∼146 nt is sufficient to initiate epigenetic silencing. This region overlaps with the 5P region of the D4Z4 (42–44), a region that is relatively hypomethylated in FSHD cells compared to controls. We show that both DNA methylation and repressive histone modifications contribute to the silencing activity of this D4Z4 region, as treatment with DNA methyltransferase inhibitor 5-aza-dC or HDAC inhibitor entinostat partly relieves its suppressive effect. In an siRNA panel of several known D4Z4 regulators, suppression of reporter expression was diminished by knockdown of SETDB1, its binding partner ATF7IP, HDACs SIN3A/SIN3B, and LRIF1. In addition, enhanced suppression of the reporter by treatment with p38 inhibitors indicates that this pathway regulates the activity of a factor, or factors, involved in epigenetic silencing, and supports the efficacy of this reporter system as a platform for FSHD drug screening.

In our assay system, only a subset of the factors previously shown to regulate D4Z4 epigenetic repression were necessary for the repressive activity of D4Z4-S5. SETDB1 and its enhancer of nuclear stability cofactor ATF7IP (46) were necessary for silencing; however, TRIM28, part of a SETDB1 silencing complex was not necessary to silence the D4Z4-S5 despite its role in maintaining silencing of the D4Z4 region as shown in several other studies (26,56–59). Future studies might test other factors in complexes with SETDB1, such as components of the HUSH complex (60–62). Importantly, genome-wide clustered regularly interspaced short palindromic repeats (CRISPR) knockout screens have the potential to discover additional factors and pathways necessary to initiate and maintain repression of D4Z4-S5.

The D4Z4-S5 DNA methylation was higher in cells with silenced GFP expression compared to GFP expressing cells. It is notable, however, that many of the silenced cells had little or no DNA methylation, indicating that silencing occurs initially through factor-mediated repression. The decreased H3K9me3 associated with knockdown of SETDB1 and ATF7IP suggest that this histone modification might be necessary to initiate silencing followed subsequently by DNA methylation. The maintenance of H3K9me3 at the D4Z4-S5 region of the endogenous D4Z4, overlapping with the 5P region, suggests that factors associated with adjacent regions might be sufficient to maintain this repressive chromatin modification.

Several screens have identified p38 inhibitors as repressors of *DUX4* expression (50,51), however, the necessary targets of this pathway have not been identified. Our study indicates that p38 pathway members may modulate expression through D4Z4-S5, though p38-inhibitor-induced silencing did not persist for the full time-course of treatment. This is consistent with a recent study showing p38 inhibition of DUX4 expression in early myogenesis but not late myogenesis (63), suggesting that compensatory pathways might limit the duration of efficacy. In this context, the D4Z4-S5 reporter cell line might be particularly useful for drug screens to identify suppressors of D4Z4 without the secondary complications of altering myogenesis and with the added value of long-term treatment monitoring. Similarly, CRISPR interference (CRISPRi) strategies might be optimized for this region.

Future studies will be needed to extend our findings and their implications for FSHD and the developmental regulation of DUX4. We used a non-muscle cell line, HeLa cells, to avoid the dynamic regulation of the D4Z4 during skeletal muscle differentiation (36) and because of the higher efficiency of integration into the safe harbor locus. Extending these studies to human skeletal muscle cells will be important to confirm that D4Z4-S5, and no other D4Z4 segments, have similar silencing activity, and to assess the response during differentiation of myoblasts to differentiated muscle cells. Based on ChIP, the enrichment of SMCHD1 and LRIF1 at the D4Z4-S5 was less substantial than at the endogenous LRIF1 promoter and, as noted above, the failure of SMCHD1 knockdown to de-repress D4Z4-S5 does not support its direct role in silencing through this region. This agrees with previous work in control myoblasts containing hypermethylated D4Z4 regions, where loss of SMCHD1 was not sufficient to alter H3K9me3 levels in the D4Z4 and led to only minor *DUX4* de-repression compared to levels present in FSHD2 myoblasts (38), and is also consistent with a recent report showing that SMCHD1 can play a role as a transcriptional co-activator in some contexts (48). Future studies will need to determine whether extending the D4Z4-S5 fragment size or creating a multi-copy array will enhance SMCHD1 repressive activity.

Ultimately, the discovery of D4Z4-S5-mediated silencing activity in this isolated reporter system provides a robust platform for identification of new D4Z4-regulating factors, pathways, and therapeutics. We used HeLa cells to establish initial proof-of-concept use of this system and believe it will be easily adaptable for other cell types. Flow cytometry of GFP expression is a simple and sensitive system for future CRISPR- or RNAi-mediated screens of epigenetic regulators that may interact with this important D4Z4-S5/5P regulatory sequence. This study also shows that cells with D4Z4-S5 can be used for temporal studies in multi-well live-cell imaging screens to detect robust changes in GFP+ cell confluency over time in response to treatment. Our hope is that this system may be freely used as a platform for future discoveries in FSHD research.

## Materials and Methods

### Cell culture

HeLa cells were grown in DMEM High Glucose with L-glutamine (Gibco, #11965092), supplemented with 10% fetal bovine serum (FBS) (Cytiva Hyclone, #SH30396.03) and 1% penicillin/streptomycin (Gibco, #15140122) and incubated at 37°C supplemented with 5% CO_2_. Cells were passaged by treatment with 0.25% trypsin-Ethylenediaminetetraacetic acid (EDTA) (Gibco, #25200056) for 5 minutes at 37°C followed by growth media addition and subculturing. For the *de novo* silencing experiment, HeLa cells were treated with growth media supplemented with 30μM 5-aza-2’-deoxycytidine (Millipore Sigma #189825), 5μM entinostat (N/A) or equivalent volume of DMSO (Sigma-Aldrich, #D2650) for 12 hours before washout and addition of growth media for 12 hours, with 12 hour on/off treatments repeated for three days. Cells were then harvested 1, 3, 7, and 10 days after the final washout for harvest of genomic DNA (gDNA) and flow cytometry analysis of GFP. Cells were treated with DUX4 inhibitors losmapimod (Sigma Aldrich, #SML3596), SB203580 (Selleckchem, #S1076), clenbuterol (Sigma Aldrich, #C5423), formoterol (Sigma-Aldrich, #F9552), (+)- JQ1 (Cayman Chemical, #11187-1), or berberine (Sigma Aldrich, #B3251) at escalating doses for three days before harvest for flow cytometry analysis.

### AAVS1 silencing reporter knock-in cell line generation

Various D4Z4, LRIF1 promoter region, and control sequences were cloned into the enhancer region of the AAVS1 HA(L)-CMV-mEmerald-hPGK-PuroR-AAVS1 HA(R), parental plasmid pMK232 (CMV-OsTIR1-PURO) Addgene#72834, which was a gift from the lab of Robert Bradley (FHCC). This plasmid was linearized by inverse PCR (Phusion Plus PCR Master Mix, Thermo Scientific #F631S), to exclude the CMV enhancer region. The backbone was purified using NucleoSpin Gel and PCR cleanup mini kit (Machery-Nagel, #740609). Inserts were amplified via PCR with primers (IDT) containing backbone-overlapping sequences with flanking HindIII and SalI cut sites for easier subsequent cloning (primers listed in Supplementary Material, Table S3) and gel purified. D4Z4 segments were amplified from the lambda42 fragment, which contains 2.5 copies of the D4Z4 (18). LRIF1 promoter region and control fragments were PCR amplified from HeLa genomic DNA. These insert fragments were cloned into the PCR-amplified or HindIII-HF-(NEB, #R3104L) and SalI-HF-(NEB, #R3138L) digested linearized AAVS1 backbone via HiFi DNA Assembly Master mix (NEB, #E2621L) according to manufacturers’ protocols. Plasmids were transformed into Stbl3 competent cells, extracted via PureLink Quick Plasmid Miniprep Kit (Invitrogen, #K210011), verified via sanger sequencing (FHCC Genomics core), and further grown and expanded for extraction via Nucleobond Xtra Maxi EF kit (Macherey-Nagel, #740424.50). AAVS1 donor plasmids and pX459-sgAAVS1 (Addgene #62988), which contains a single guide RNA (sgRNA) for the AAVS1 site and Cas9, were transfected into HeLa cells with Lipofectamine 3000 (Invitrogen, #L3000015) according to manufacturer’s protocol. HeLa lines were selected for proper AAVS1 knock-in using puromycin selection (Sigma-Aldrich, #P8833) at 1.5μg/mL. Cells were cultured in puromycin for 1 week and then sorted for GFP^High^ cells via FACS. These GFP+ lines were then expanded without puromycin for two weeks before experimentation and analysis for GFP expression levels via flow cytometry. All cloning and sequencing primers are listed in Supplementary Material, Table S3.

### Flow cytometry and FACS

Cells were trypsinized and resuspended in FACS Buffer (1X phosphate buffered saline (PBS) with 2%FBS and 5mM EDTA) and kept on ice until analysis. For flow cytometry analysis, live cell samples were run on one of several analyzers (BD Biosciences: FACSCanto II, LSR Fortessa x50, FACSymphony A3, or FACSymphony A5). For FACS, cells were sorted into tubes or plates with growth media using a BD Biosciences FACSymphony S6 or FACS Aria II instrument. GFP (mEmerald) fluorescence was quantified using Blue 488nm (200mW) laser readouts. For all samples shown on the same graph, the same voltage parameters and machine was used for consistency. This analysis was supported by the Flow Cytometry Shared Resource (RRID:SCR_022613) of the Fred Hutch/University of Washington/Seattle Children’s Cancer Consortium (P30 CA015704). FlowJo v10.8 software (BD Biosciences) was used for all flow cytometry and FACS analysis. For all GFP fluorescence intensity graphs, cells were first gated for live cells to eliminate debris using forward and side scatter (FSC and SSC), then gated on single cells to eliminate doublets using FSC-Height and FSC-Width. GFP channel values reflect log scale distribution of fluorescence intensity.

### Quantitative reverse transcription PCR (RT-qPCR)

Total cellular RNA was isolated using the Nucleospin RNA kit (Machery-Nagel, #740955.250) according to the manufacturer protocol. 0.5-1μg RNA was treated with Amplification grade-DNaseI (Thermo Fisher, #18068015), heat inactivated with EDTA (Thermo Fisher, #AM9260G), and reverse transcribed into cDNA using random hexamers or oligo-dT with the Superscript IV First-Strand Synthesis System (Invitrogen, #18091050) following manufacturer instructions. Quantitative PCR was run on a QuantStudio 7 Flex (Applied Biosystem) using iTaq SYBR Green Supermix (Bio-Rad, #1725124). Primers are listed in Supplementary Material, Table S3.

### Immunoblotting

Cells were lysed in radioimmunoprecipitation assay (RIPA) buffer (150mM NaCl, 1% NP-40, 0.5% Na-deoxycholate, 1% sodium dodecyl sulfate (SDS), 25mM Tris-HCl pH7.4) with protease and phosphatase inhibitors (Pierce, #A32961) Protein lysate was sonicated in a Bioruptor Sonication System (Diagenode) and cleared by centrifugation at 16,000xg. Lysate was quantified via Pierce bicinchoninic acid (BCA) assay (Thermo Scientific, #23225). Protein lysates were run on NuPAGE 4-12% Bis Tris polyacrylamide gels (Invitrogen, #NP0322BOX) using NuPAGE MOPS buffer (Invitrogen, #NP0001) and transferred onto polyvinylidene fluoride (PVDF) membranes (Invitrogen, #LC2002). Membranes were blocked in PBS-Tween (PBST) buffer (0.1% Tween-20 in 1XPBS) with 5% non-fat dry milk before incubation overnight at 4°C with primary antibody diluted in PBST buffer (primary antibodies listed in Supplementary Material, Table S3). Membranes were incubated in horseradish peroxidase-conjugated secondary antibody (Supplementary Material, Table S3) for one hour at room temperature. SuperSignal chemiluminescent substrate (Thermo Scientific, #34580) was added to membranes and a Mini-medical 90 processor (AFP Manufacturing) was used for detection on film.

### siRNA transfections

HeLa AAVS1 reporter cells were seeded into 6-well plates 24 hours before transfection. 25-50pmol of siRNA was diluted in OptiMEM (Gibco, #31985070) and transfected into cells using Lipofectamine RNAiMAX (Invitrogen #13778150) following manufacturer’s instructions. 48 hours after transfection, wells were changed to fresh growth media and the siRNA transfection was repeated. 48 hours after the second transfection, cells were harvested for flow cytometry of GFP fluorescence and RNA isolation for RT-qPCR analysis.

### Bisulfite sequencing and % CpG Methylation Analysis

HeLa AAVS1 reporter lines were harvested, pelleted, and flash frozen in a dry ice/ethanol slurry before storage at -80°C. Genomic DNA was isolated using the GeneJet Genomic DNA Purification kit (Thermo Scientific, #K0722) following manufacturer instructions. 200-500ng of genomic DNA was subjected to bisulfite conversion using EZ DNA Methylation-Gold Kit (Zymo Research, #D5005) following manufacturer instructions. Bisulfite-converted DNA was then used for PCR amplification of the AAVS1 insert regions using ZymoTaq DNA Polymerase (Zymo Research, #E2001) and degenerate primers (IDT, Supplementary Material, Table S3). PCR products were gel purified using the NucleoSpin Gel and PCR cleanup mini kit (Machery-Nagel, #740609) and sequenced using Premium PCR Sequencing performed by Plasmidsaurus using Oxford Nanopore Technology with custom analysis and annotation. Reads were aligned to insert contigs using BWAMeth.py (64) then filtered to only includes reads covering 90% of the contig length using samtools (65). After subsetting, reads were imported into R (66) using GenomicAlignments (67), converted to a dataframe via BioStrings (68) and percent methylation was calculated using the tidyverse package of tools (69). Raw sequencing reads are presented in Supplementary Material, Table S2.

### ChIP-qPCR

ChIP-qPCR was adapted from a previously published protocol (70). HeLa AAVS1 cells were trypsinized, washed with 1xPBS, and fixed in 1% Formaldehyde Ultrapure EM grade (EMS, #15710) for 10 minutes at room temperature with gentle rocking. Formaldehyde was quenched with 125mM glycine, cells were washed with PBS and then lysed in Lysis Buffer (Protease and phosphatase inhibitors (Pierce, #A32961), 1% Na-deoxycholate, 1% SDS, 5mM EDTA pH 8, and 50mM Tris pH 8 in 1xPBS). Lysate was sonicated in Biorupter Plus TPX Polymethylpentene tubes (Diagenode, C30010010) using a Biorupter Plus Sonication System (Diagenode, B01020001) until chromatin was sheared to a range of 200-600bp; a small aliquot of sheared chromatin was incubated with RNase A at 37°C for one hour, treated with Proteinase K overnight at 65°C, and purified using a NucleoSpin Gel and PCR cleanup mini kit (Machery-Nagel, #740609) before concentration was determined using a Qubit and chromatin was run on a 2% agarose gel in Tris/Borate/EDTA (TBE) buffer to assess fragment size. Chromatin was centrifuged at 16,000xg for 10 min at 4°C to clear insoluble fraction, and 10% of the chromatin was reserved for input DNA. For immunoprecipitation, chromatin was diluted in Dilution Buffer (1% Triton X-100, 150mM NaCl, 2mM EDTA pH 8, and 20mM Tris pH 8 in PBS) to reduce SDS concentration to <0.1%. For LRIF1 and SMCHD1 ChIP, 5μg of antibody (rbαSMCHD1 Abcam #Ab31865, rbαAnti-Ligand-dependent nuclear receptor-interacting factor 1, Millipore #ABE1008, or Rabbit IgG control Millipore #PP64) was added to 30μg of chromatin and incubated with rotation overnight at 4°C. Agarose Fast Flow Prot A (Millipore Sigma #16-156) beads were pre-blocked overnight at 4°C in Dilution Buffer + 2% bovine serum albumin (BSA). For histone ChIP, 2-4μg of antibody (rbαH3K9me3 Abcam #Ab8898, or Rabbit IgG control Millipore #PP64) was added to ∼2-5μg of chromatin and incubated with rotation overnight at 4°C. Dynabeads Protein A (Invitrogen #10002D) beads were pre-blocked overnight at 4°C in Dilution Buffer + 2% BSA. Samples were incubated with beads at 4°C for 2 hours, and washed with Wash Buffer I (protease and phosphatase inhibitors (Pierce, #A32961), 0.1% SDS, 1% Triton X-100, 2mM EDTA pH 8, 20mM Tris pH 8, 150mM NaCl), washed with Wash Buffer II (protease and phosphatase inhibitors, 0.1% SDS, 1% Triton X-100, 2mM EDTA pH 8, 20mM Tris pH 8, 500mM NaCl), washed with Wash Buffer III (protease and phosphatase inhibitors, 0.25M LiCl, 1% NP-40, 1% Na-deoxycholate, 1mM EDTA pH 8, 10mM Tris pH 8, 500mM NaCl), and washed twice with ice cold PBS. Beads and attached chromatin were then de-crosslinked by resuspension in Ultrapure water with 1% SDS and 0.1M NaHCO3, incubated with RNase A (Thermo Scientific, # R1253) for 1 hour at 37°C, and incubated overnight at 65°C with Proteinase K (Thermo Scientific, #EO0491). IP DNA was then purified using NTB Buffer (Machery-Nagel, #740595.150) and the NucleoSpin Gel and PCR cleanup mini kit (Machery-Nagel, #740609). Quantitative PCR of IP DNA was run on a QuantStudio 7 Flex (Applied Biosystem) using iTaq SYBR Green Supermix (Bio-Rad, #1725124). Primers are listed in Supplementary Material, Table S3.

### Incucyte tracking of GFP+ cell confluence

HeLa AAVS1 lines were seeded at 80% confluency and 24 hours later were treated with 5-aza-2’-deoxycytidine (Millipore Sigma #189825) for 12 hours. Cells were then incubated with standard media for 12 hours; this was repeated twice more, for a total of 36 hours of 5-aza-2’-deoxycytidine treatment in 72 hours. 24 hours after washing out the third round of 5-aza-2’-deoxycytidine and replacement with standard media, GFP+ cells were sorted by FACS at a density of 10,000 cells per well in a 96-well plate with 100uL media per well. 24 hours later, 100uL of media with 2x concentration of losmapimod (Sigma Aldrich, SML3596) or dorapimod (MedChem Express, #HY-10320) (1μM, final concentration 0.5μM) and 96-well plates were placed in the Incucyte S3 Live Cell Analysis system, kept at 37°C with 5% CO2. Wells were imaged for GFP+ and brightfield cell confluency, cell count, and GFP Mean Fluorescence intensity every 2 hours for 6 days. On day 3, plates were removed from the Incucyte between images and media was changed and supplemented with fresh p38 inhibitors. GFP+ cell confluency per well was normalized to t0 for that well and to the average of DMSO treated wells to analyze differences in p38 inhibitor-treated wells.

## Acknowledgments

We thank Dr. Robert Bradley (FHCC) for the AAVS1 HA(L)-CMV-mEmerald-hPGK-PuroR-AAVS1 HA(R) plasmid. This research was supported by the Flow Cytometry Shared Resource (RRID:SCR_022613) and the Genomics and Bioinformatics Shared Resource (RRID:SCR_022606) of the Fred Hutch/University of Washington/Seattle Children’s Cancer Consortium (P30 CA015704). We thank Dr. Patrick Paddison, Dr. Adam Geballe, Dr. Andrew Hsieh, Dr. Sita Kugel, and Dr. Toshio Tsukiyama (FHCC) for their perspectives regarding this project. We also thank members of the Tapscott lab for discussion and critical reading of the manuscript.

## Supplementary Material

**Figure S1:**
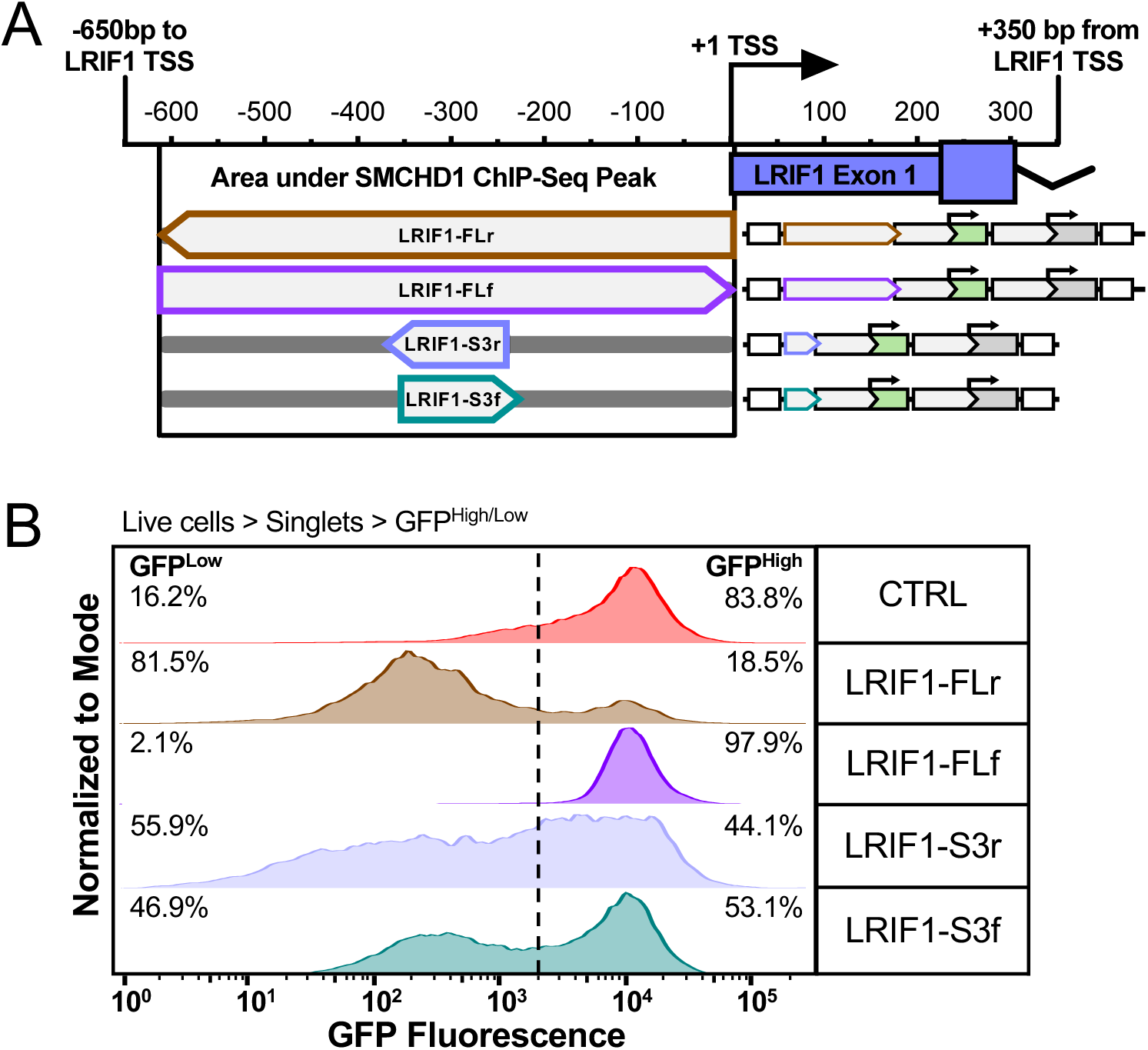
Silencing activity of LRIF1-S3, but not LRIF1-FL, is independent of sequence orientation. **(A)** Schematic of LRIF1-FLr, LRIF1-FLf, LRIF1-S3r and LRIF1-S3f orientation relative to the genomic sequence and the integrated silencing reporter construct. **(B)** Flow cytometry GFP fluorescence histogram of LRIF1-FLr/f and LRIF1-S3r/f compared to CTRL cells, normalized to mode.

**Figure S2:**
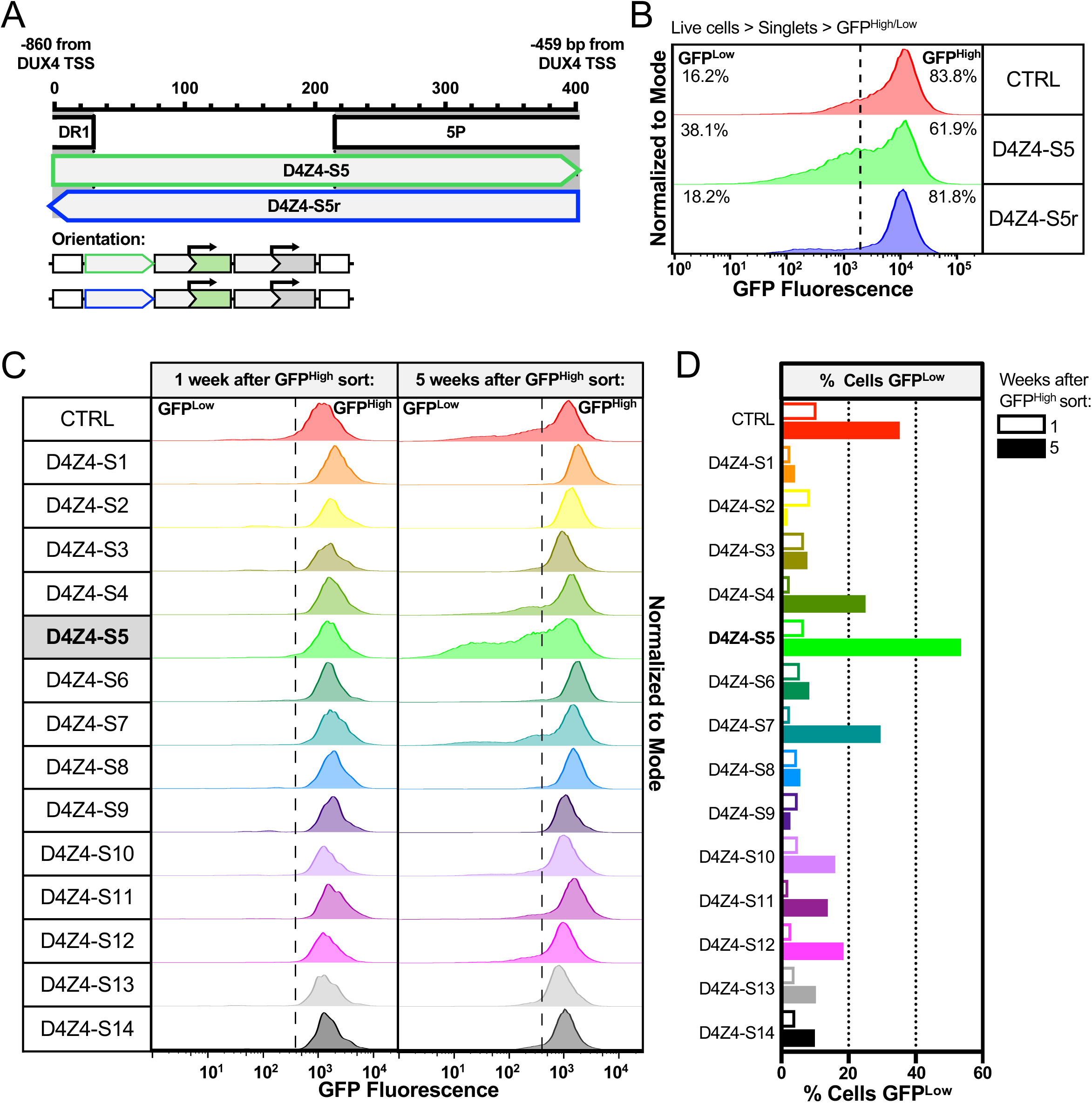
D4Z4-S5 silencing activity is orientation independent, transient, and increases over time in culture. **(A)** Schematic of D4Z4-S5 and D4Z4-S5r orientation relative to the genomic sequence and the integrated silencing reporter construct. **(B)** Flow cytometry GFP fluorescence histogram of D4Z4-S5 and D4Z4-S5r compared to CTRL cells, normalized to mode. **(C)** Flow cytometry GFP fluorescence histogram of D4Z4 silencing construct segments 1 (left) and 5 (right) weeks after GFP^High^ FACS sort, normalized to mode. **(D)** Percentage of GFP^Low^ cells in each D4Z4 silencing construct line 1 (clear) or 5 (filled) weeks after GFP^High^ FACS sort.

**Figure S3:**
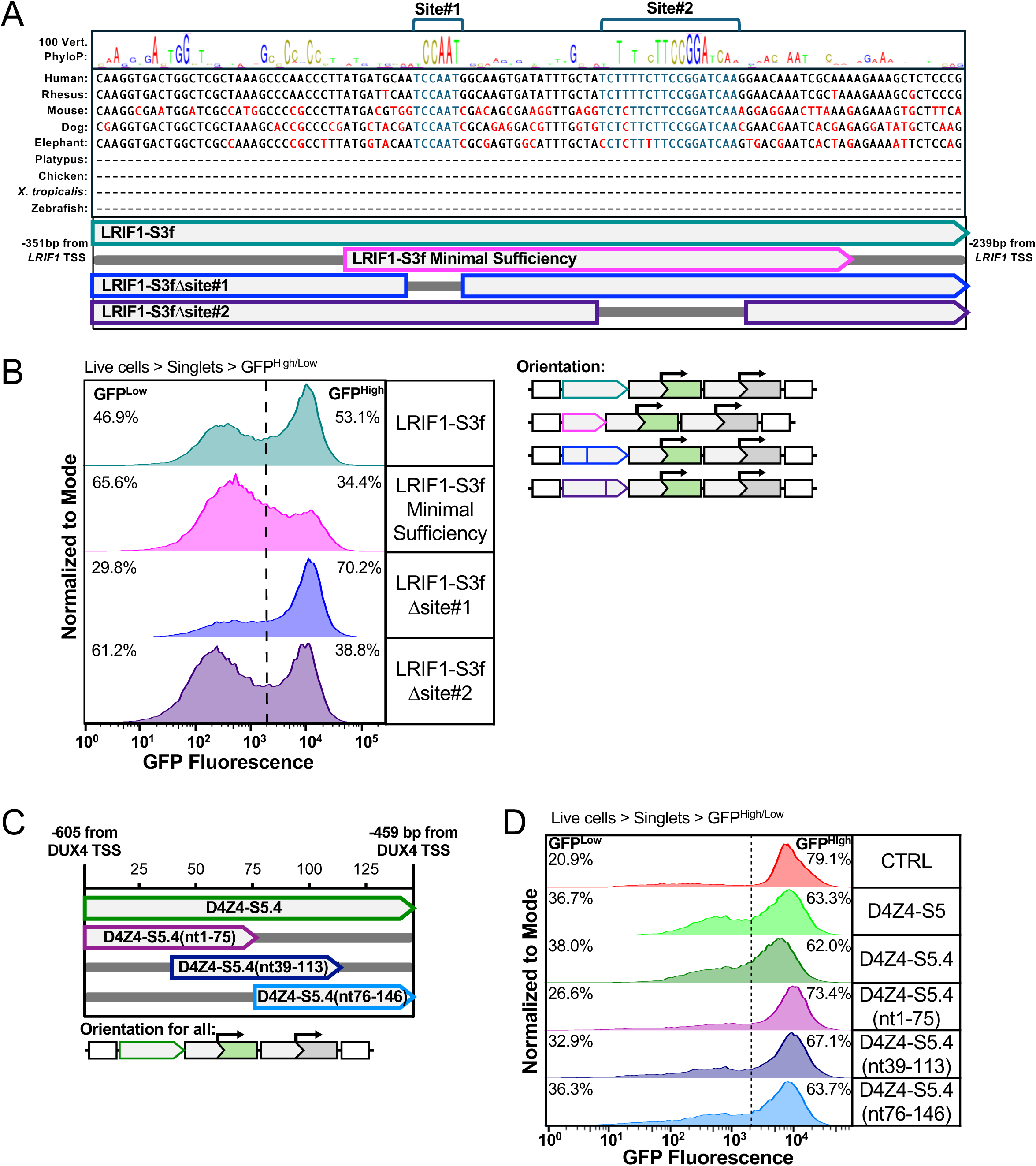
LRIF1-S3, and not D4Z4-S5.4, contains a conserved sequence that conveys silencing activity. **(A)** Top: Logo plot of LRIF1-S3 basewise conservation consensus sequence based on UCSC genome browser phyloP analysis of 100 vertebrates. Middle: LRIF1-S3 sequences in 9 representative species show conservation only in placental mammals. Bottom: Schematic of LRIF1-S3 conserved site deletion inserts. **(B)** Flow cytometry GFP fluorescence histogram of LRIF1-S3 conserved site deletion inserts, normalized to mode. Percentages of cell populations classified GFP^High^ or GFP^Low^ are indicated. **(C)** Schematic of D4Z4-S5.4 sub-fragment inserts. **(D)** Flow cytometry GFP fluorescence histogram of D4Z4-S5.4 sub-fragment inserts, normalized to mode. Percentages of cell populations classified GFP^High^ or GFP^Low^ are indicated.

**Figure S4:**
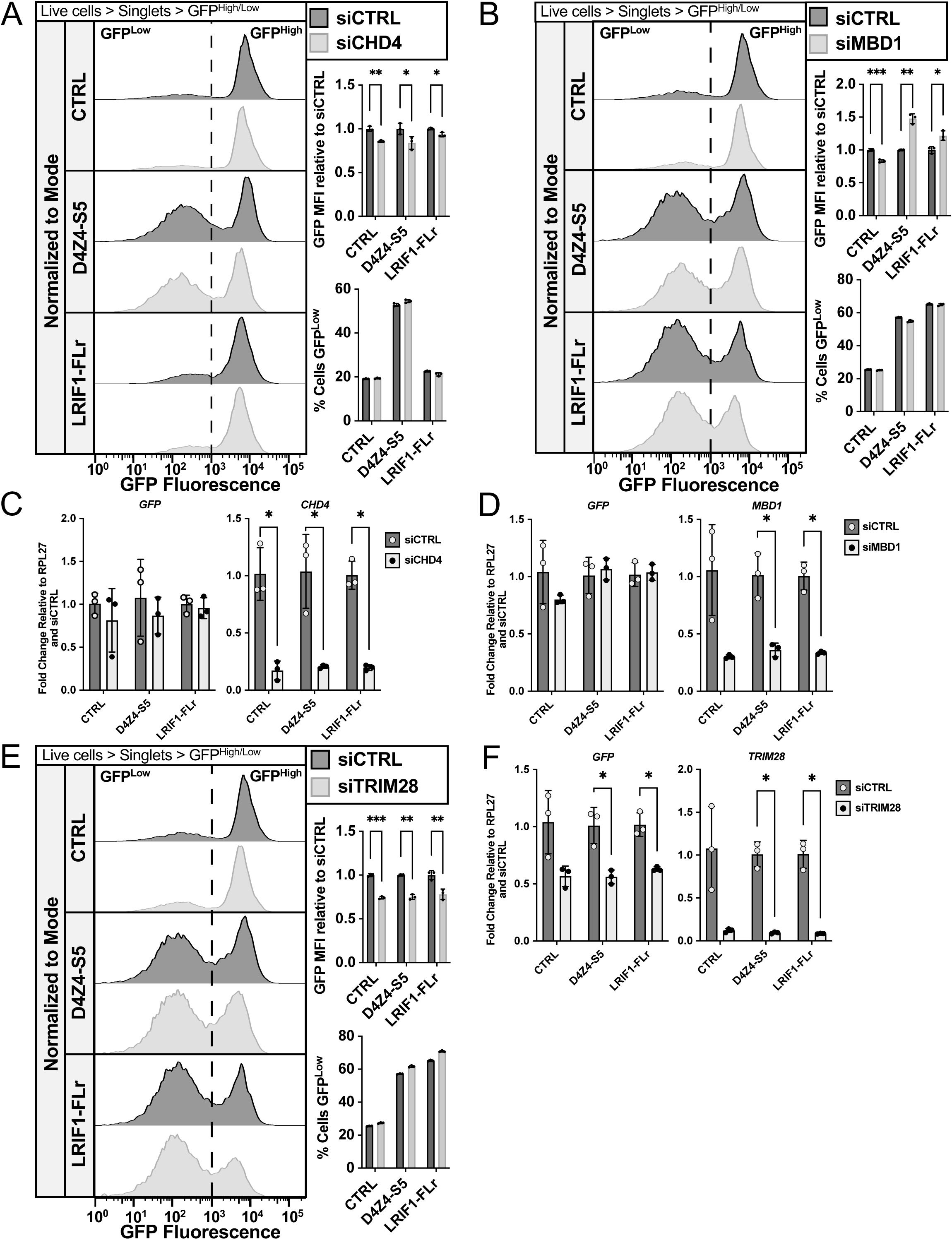
Knockdown of MBD1, CHD4, or TRIM28 does not rescue GFP expression in D4Z4-S5 and LRIF1-FLr cells. **(A-B, E)** Left: Representative singleton GFP fluorescence histograms of CTRL, D4Z4-S5 and LRIF1-FLr cells treated with CTRL or CHD4 **(A)**, MBD1 **(B)**, or TRIM28 **(E)** siRNAs. Histograms are normalized to the mode of the population. Right Top: Fold change in GFP MFI upon siRNA-mediated knockdown of CHD4 **(A)**, MBD1 **(B)**, or TRIM28 **(E)** compared to siCTRL. Data represent mean ± SD of biological replicates, n=3. Statistical significance was determined by Welch’s t-test: *p<0.05, **p<0.01, ***p<0.001. Right bottom: Percentage of GFP^Low^ cells in the population at various time points after siRNA treatment. Data represent mean ± SD of biological replicates, n=3. **(C-D, F)** RT-qPCR analysis of *GFP* (left) and *CHD4* **(C)**, *MBD1* **(D)** or *TRIM28* **(F)** (right) expression compared to housekeeping gene *RPL27* expression in siRNA-treated CTRL, D4Z4-S5, and LRIF1-FLr cells. Data plotted is fold change of expression in siCHD4-(**C)**, siMBD1-**(D)**, or siTRIM28-**(F)** treated cells compared to siCTRL-treated cells. Data represent mean fold change ± SD of biological replicates, n=3. Statistical significance was determined by Welch’s t-test: *p<0.05.

**Figure S5:**
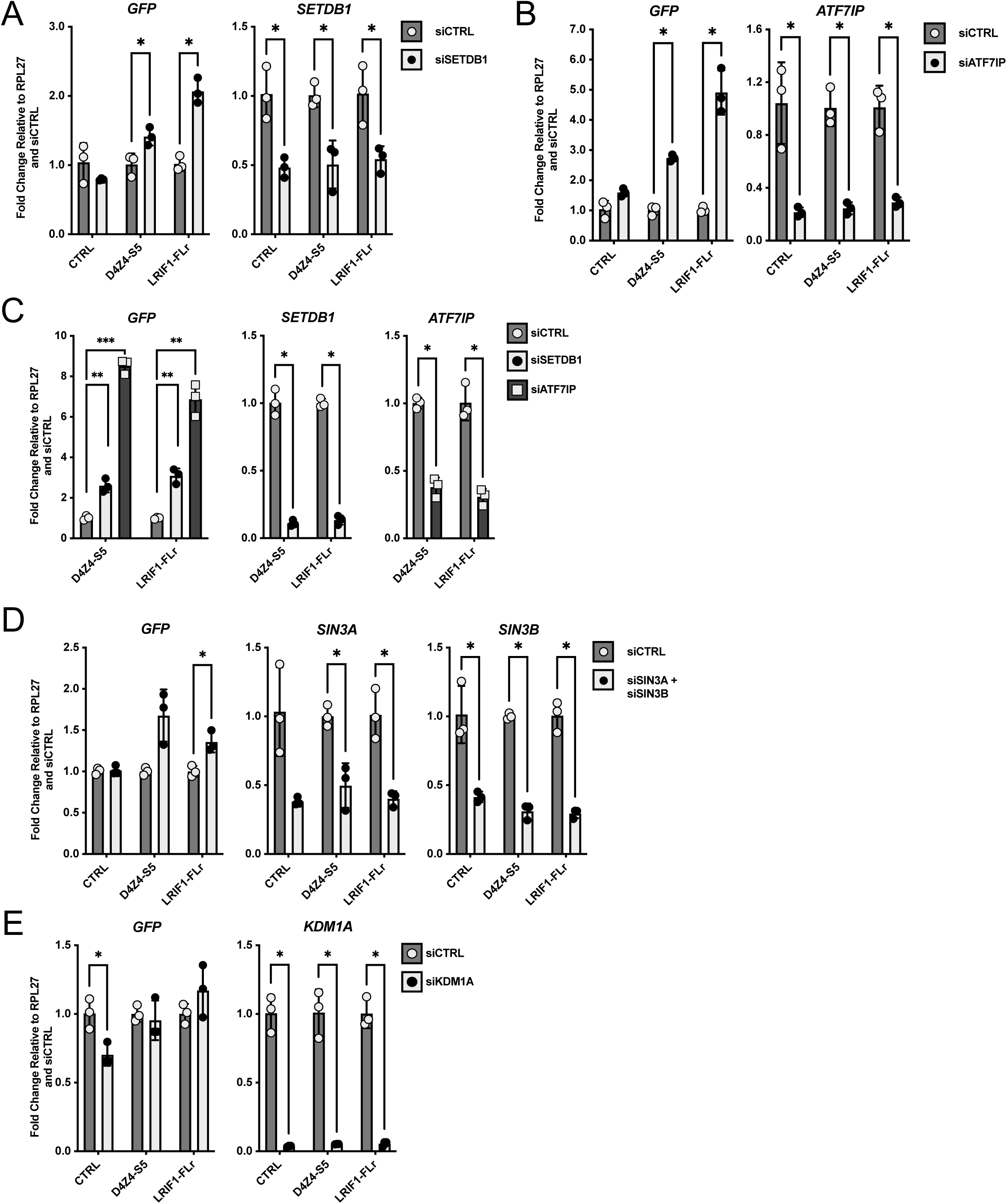
RT-qPCR analysis confirms GFP expression changes and sufficient knockdown efficiency of identified epigenetic regulators. **(A-E)** RT-qPCR analysis of *GFP* (left) and *SETDB1* (A), *ATF7IP* (B), *SETDB1 and ATF7IP* from H3K9me3 ChIP samples (C), *SIN3A* and *SIN3B* (D), or *KDM1A* (E) (right) expression compared to housekeeping gene *RPL27* expression in siRNA-treated CTRL, D4Z4-S5, and LRIF1-FLr cells. Data plotted is fold change of expression in siSETDB1-**(A,C)**, siATF7IP-**(B,C)**, siSIN3A/B-**(D)**, or siKDM1A-**(E)** treated cells compared to siCTRL-treated cells. Data represent mean fold change ± SD of biological replicates, n=3. Statistical significance was determined by Welch’s t-test: *p<0.05, **p<0.01, ***p<0.001.

**Figure S6:**
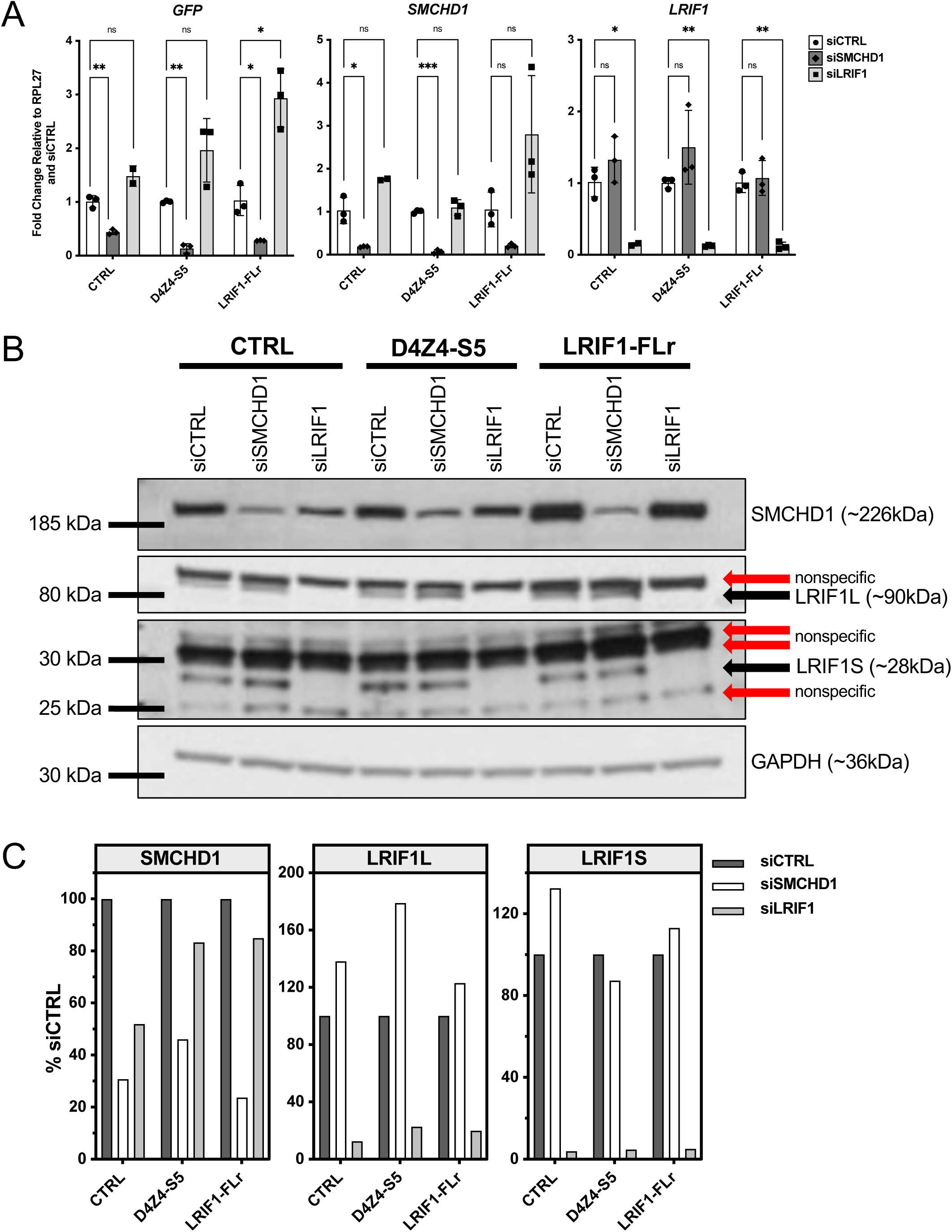
RT-qPCR and Western blotting analyses confirm changes in GFP expression and sufficient knockdown of SMCHD1 and LRIF1. **(A)** RT-qPCR analysis of *GFP* (left), *SMCHD1* (middle) and *LRIF1* (right) expression compared to housekeeping gene *RPL27* expression in siRNA-treated CTRL, D4Z4-S5, and LRIF1-FLr cells. Data plotted is fold change of expression in siSMCHD1- or siLRIF1-treated cells compared to siCTRL-treated cells. Data represent mean fold change ± SD of biological replicates, n=3. Statistical significance was determined by Welch’s t-test: *p<0.05, **p<0.01, ***p<0.001. **(B)** Immunoblot analysis of SMCHD1 and LRIF1 protein levels in CTRL, D4Z4-S5, and LRIF1-FLr cells treated with siCTRL, siSMCHD1, or siLRIF1. GAPDH serves as a loading control. **(C)** Quantification of SMCHD1 (left), LRIF1L (middle), and LRIF1 (right) levels in (B) using densitometry analysis normalized to GAPDH and graphed as percentage relative to siCTRL-treated samples.

**Figure S7:**
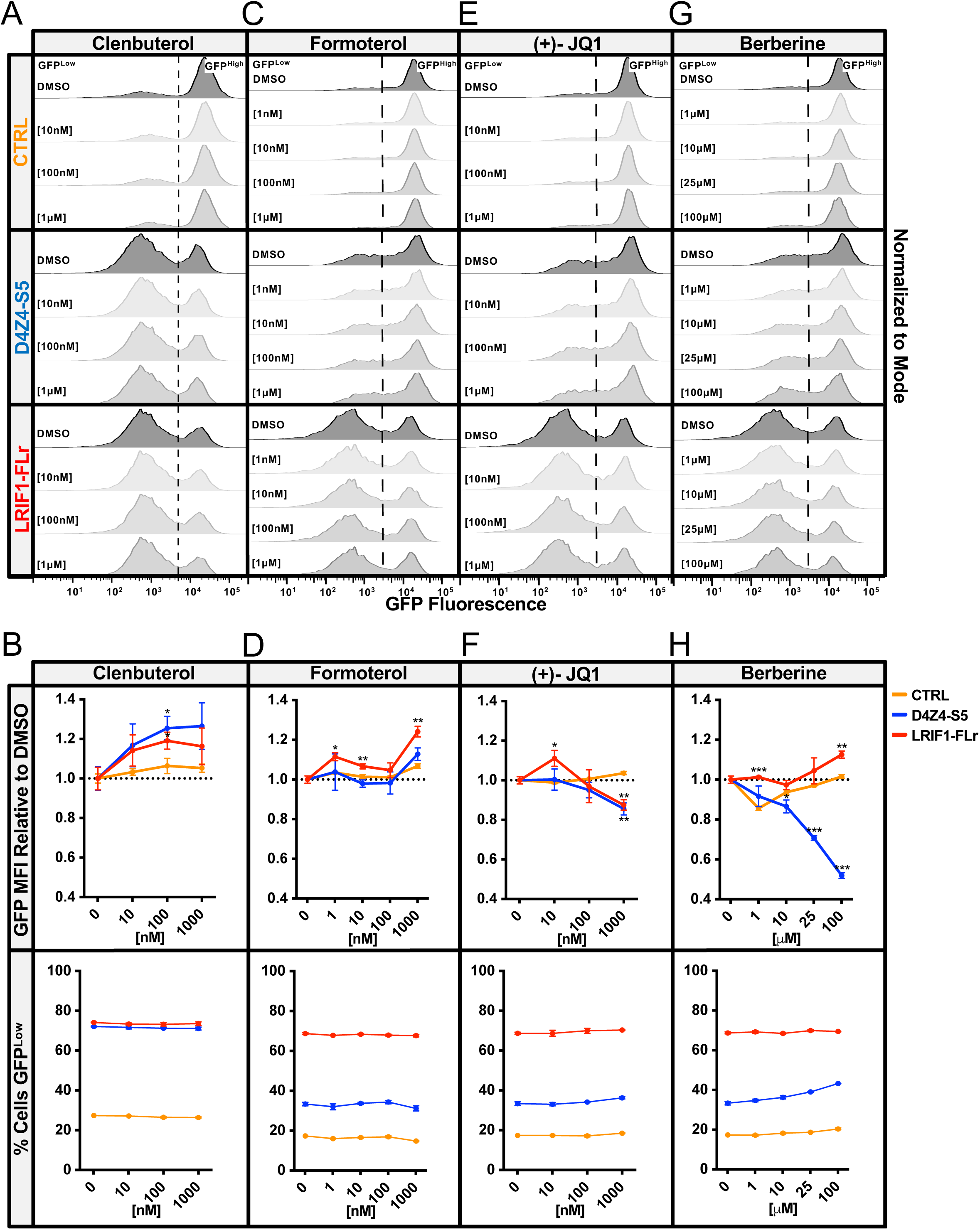
D4Z4-S5 and LRIF1-FLr show differential sensitivity to candidate FSHD therapeutics. **(A, C, E, G)** Representative singleton GFP fluorescence histograms of CTRL, D4Z4-S5 and LRIF1-FLr cells treated for 3 days with DMSO or escalating dosages of candidate FSHD therapeutics clenbuterol **(A)**, Formoterol **(C)**, (+)- JQ1 **(E)**, or Berberine **(G)**. Histograms are normalized to the mode of the population. **(B, D, F, H)** Top: Fold change in GFP MFI upon 3-day treatment with escalating dosages of clenbuterol **(B)**, Formoterol **(D)**, (+)- JQ1 **(F)**, or Berberine **(H)** compared to DMSO treatment in CTRL (orange), D4Z4-S5 (blue), and LRIF1-FLr (red) cells. Data points represent mean fold change ± SD of biological replicates, n=3. Statistical significance was determined by Welch’s t-test: *p<0.05, **p<0.01, ***p<0.001. Bottom: Percentage of GFP^Low^ cells in the population upon treatment with escalating dosages of clenbuterol **(B)**, Formoterol **(D)**, (+)- JQ1 **(F)**, or Berberine **(H)**. Data represent mean ± SD of biological replicates, n=3.

**Table S1: Length, GC, and CpG content of silencing construct inserts.**

**Table S2: Individual CpG methylation status based on the average across all sequencing reads.**

**Table S3: Oligonucleotides, constructs, reagents and resources used in this study.**

## Abbreviations

FSHD: facioscapulohumeral dystrophy
DUX4: double homeobox 4
kb: kilobase
SMCHD1: structural maintenance of chromosomes flexible hinge domain containing 1
LRIF1: ligand-dependent nuclear receptor interacting factor 1
DNMT3B: DNA methyltransferase 3B
NuRD: nucleosome remodeling and deacetylase
CAF-1: chromatin assembly factor 1
HDAC: histone deacetylase
SIN3A/3B: SIN3 transcriptional regulator family member A and B
H3K9me3: histone H3 lysine 9 trimethylation
HKMT: histone lysine methyltransferase
SETDB1: SET domain bifurcated histone lysine methyltransferase 1
TRIM28: tripartite motif containing 28
SUV39H1: suppressor of variegation 3–9 homolog 1
RUVBL1: RuvB like AAA ATPase 1
KDM1A: lysine demethylase 1A
YY1: Yin Yang 1
RNAi: RNA interference
DICER1: dicer 1, ribonuclease III
AGO2: argonaute RISC catalytic component 2
HP1γ: heterochromatin protein 1 gamma
CTCF: CCCTC-binding factor
ChIP: chromatin immunoprecipitation
BET: bromodomain and extraterminal
siRNA: small interfering RNA
ATF7IP: activating transcription factor 7 interacting protein
GFP: green fluorescence protein
nt: nucleotide
bp: base pair
TSS: transcriptional start site
PCR: polymerase chain reaction
FACS: fluorescence activated cell sorting
5-aza-dC: 5-Aza-2′-Deoxycytidine
KD: knockdown
MFI: median fluorescence intensity
qPCR: quantitative polymerase chain reaction
GDR: gene desert region
RPL13A: ribosomal protein L13a
H3K4: histone H3 lysine 4
DMSO: dimethyl sulfoxide
CRISPR: clustered regularly interspaced short palindromic repeats
CRISPRi: CRISPR interference
FBS: fetal bovine serum
EDTA: ethylenediaminetetraacetic acid
gDNA: genomic DNA
sgRNA: single guide RNA
PBS: phosphate buffered saline
FSC: forward scatter
SSC: side scatter
RT-qPCR: quantitative reverse transcription polymerase chain reaction
RIPA: radioimmunoprecipitation assay
SDS: sodium dodecyl sulfate
BCA: bicinchoninic acid
PVDF: polyvinylidene fluoride
PBST: phosphate buffered saline Tween
TBE: tris borate EDTA

